# Deep-learning predictions of biomolecular structures : persistent limitations and new horizons extended by explicit ion addition

**DOI:** 10.64898/2026.07.24.740587

**Authors:** Jules Marien, Sujith Sritharan, Beatrice Caviglia, Raphaelle Versini, Lucas Auclair, Pierre Barraud, Carine Tisné, Alaa Reguei, Samuel Murail, Fabrice Leclerc, Elise Duboué Dijon, JianJun Tao, Nathalie Basdevant, Marc Baaden, Chantal Prévost, Sophie Sacquin-Mora, Antoine Taly

## Abstract

The advent of deep learning-driven tools such as AlphaFold has revolutionized the prediction of biomolecular structures, offering unprecedented accuracy and accessibility for proteins, RNA, and their complexes. While these tools have demonstrated remarkable success in benchmarking competitions and enabled experimentalists to generate models with ease, their widespread use has also highlighted persistent challenges. These include difficulties in assessing model confidence, limitations in predicting transmembrane domains, nucleic acids, conformational diversity, and interactions with ions or ligands, as well as the tendency to misfold intrinsically disordered regions (IDRs). In this perspective, we critically evaluate the strengths and limitations of current AI-based structure prediction tools through illustrative examples with a particular emphasis on the impact of explicit ion modelling. We notably report how the explicit addition of a few potassium cations to the prediction of IDRs or G-quadruplexes can trigger massive conformational switches compared to “dry” predictions. On this basis, we suggest modelling sequences both “dry” and in the presence of explicit potassium cations as a simple, practical way to sample alternative conformations and to expose disordered regions that current predictors tend to over-fold. We discuss the importance of reporting confidence metrics in publications to avoid overinterpretation. Furthermore, we address the unique challenges of RNA structure prediction, where data scarcity and structural complexity limit the performance of both classical and deep learning methods. Our analysis underscores the need for continued methodological advancements, integration of complementary computational tools, and expansion of high-quality experimental datasets.

**TOC Graphic:** 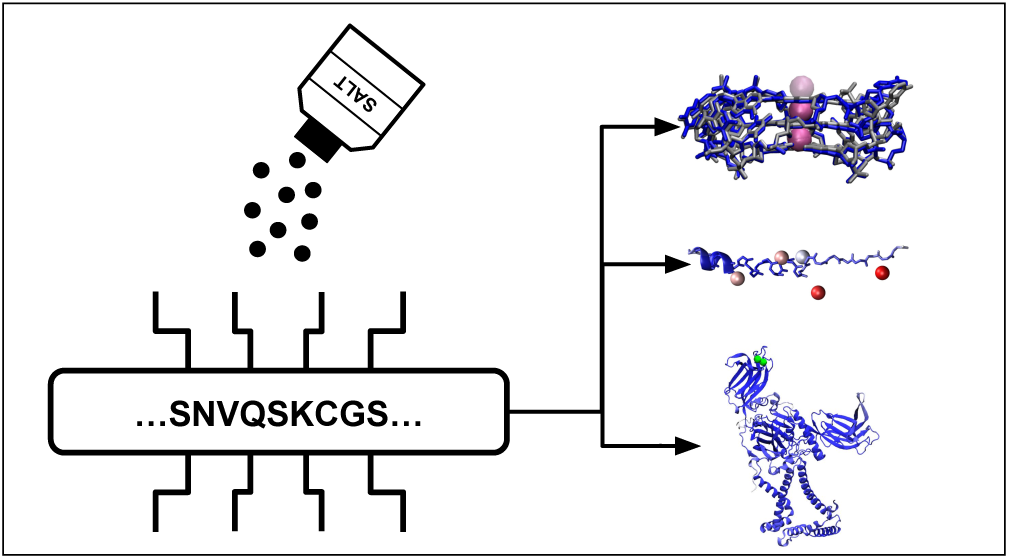

## INTRODUCTION

The past decade has witnessed a paradigm shift in structural biology, driven by the advent of deep learning–based tools for biomolecular structure prediction. The release of AlphaFold2 (AF2) and RoseTTAFold (RF) marked a turning point, achieving near-experimental accuracy in protein structure prediction, as demonstrated by their impressive performance at CASP14 and CASP15.^1^ These breakthroughs encouraged their use by structural biologists and the creation of alternatives like Boltz1, OmegaFold, and ESMFold, underscoring their transformative impact on the field. ^2,3^ The subsequent release of AlphaFold3 (AF3) further expanded the scope of prediction, enabling the modeling of not only proteins but also nucleic acids, small molecules and ions.^4,5^ These accomplishments were ultimately recognized with the 2024 Nobel Prize in Chemistry.

These advances have also facilitated the construction of biologically representative systems, thanks to the integration of AI-driven predictions with experimental data. For example, it has made possible the prediction of complex assemblies such as the NOX system^6^ and the recent integrative modeling of mitochondrial cristae, which combined molecular dynamics with coarse-grained simulations to capture the complexity of cellular environments. ^7^ Constructing models of those systems was considered beyond reach only a few years ago and is blurring the traditional split between molecular and cell biology.^8^

However, significant challenges remain. While tools like AF3 and RFNA have made it remarkably easy for non-specialists to generate structural models, assessing their quality and reliability can be difficult, especially in prospective settings. Predictive programs often excel in retrospective validation but may struggle with conformational diversity, allosteric transitions, protein–ligand cofolding, and the dynamic ensembles that underpin biological function.^9–11^

This perspective explores with selected examples the impact of these revolutionary tools on structural biology, their integration with experimental approaches, and the remaining hurdles that must be addressed to fully realize their potential in both fundamental research and applied sciences.

We first assess the new modelling possibilities achieved by the current deep-learning tools, such as predictions of nucleic acids, transmembrane proteins and ligands, before discussing their main limitations. A particular focus is made on the capacity of new models such as AF3 to explicitly model ions and capture ion-induced structural changes. We show with specific examples that, while still far away from accuracy, this new feature of predictors can be hijacked to enhance and diversify their capabilities. We conclude with a discussion on model assessment, exposing available metrics and tools to critically consider the validity of a prediction.

### New modeling horizons

Deep-learning structural prediction tools for biomolecular systems first demonstrated their accuracy for folded cytoplasmic proteins. Since the emergence of AlphaFold, new prediction horizons are now being explored such as the modeling of transmembrane domains, nucleic acids and explicit ions. We first use RNA modeling as an opportunity to explain recent advances in deep-learning prediction schemes, exemplifying the notions with the TrmK–tRNA complex. We next develop the cases of mitofusins and MCTP proteins as archetypal examples of the challenges and successes encountered using deep-learning predictions of their transmembrane domains. MCTPs are suspected to bind calcium cations, ^12^ and we expose the different tools and strategies available to explicitly predict this interaction. We conclude this part with an original analysis of the adenylate kinase (Adk) protein family, and how explicit ion modelling can trigger the sampling of different conformations by AF3.

### I. RNA structure prediction

Predicting the three-dimensional (3D) structures of ribonucleic acids (RNA) remains a central challenge in computational biology because RNA function is intimately tied to its 3D conformation. RNA participates in catalysis, gene expression and regulation, and to a host of other cellular processes. Hence, many emerging therapeutic strategies target RNA directly. Experimental structure determination by X-ray crystallography, cryo-electron microscopy (cryo-EM) or NMR is still labor-intensive, costly, and often hampered by the intrinsic flexibility of RNA. Consequently, computational modelling has become an indispensable complement to experimental work.^13^

Over the past two decades, a variety of non-machine-learning (non-ML) methods such as fragment-assembly, physics-based simulations and knowledge-based potentials have been developed.^14^ More recently, deep-learning (DL) approaches that generate structures from the sequence alone have shown remarkable success for proteins (e.g., AlphaFold2) and are now being adapted to RNA.^15^ The dramatic gains observed for proteins stem largely from the abundance and high quality of protein structures in the Protein Data Bank (PDB). By contrast, RNA suffers from several data-related bottlenecks that limit the performance of both classical and DL methods.^14^

### I.A. From secondary structure to motif prediction

Deep neural networks have already surpassed traditional algorithms for RNA secondary-structure inference and motif detection,^16^ and benchmark studies report that modern ML methods generally achieve higher accuracy than classical pipelines. ^17^ Nevertheless, the advantage narrows dramatically for newer folds not included in the AI training set or synthetic RNAs, as these successes rely on training and test sets that share similar sequence and structural distributions^18^. Performance is strongly modulated by three factors:

1. **RNA family and evolutionary depth:** Methods that depend on deep multiple-sequence alignments (MSAs), such as RoseTTAFold-RNA, perform poorly on ‘orphan’ RNAs where MSA depth is low^17^.
2. **Sequence length and fold complexity:** Double-helical and cloverleaf motifs, which dominate the PDB, are predicted with higher fidelity than intricate tertiary folds^19^.
3. **Similarity to training data:** Across several studies, the Pearson correlation between predicted TM-score and the maximal structural similarity to any training exemplar ranges from 0.72 (AlphaFold-3 RNA^19^) to 0.78 for RNA-protein complexes^19^. When similarity falls below a TM-score of 0.25, the mean predicted TM-score drops below 0.1, indicating near-random performance.

AlphaFold-3 and Boltz-1 are currently at the forefront of deep-learning frameworks for RNA structure prediction. For isolated RNA chains, the two methods deliver comparable accuracy, but AF3 gains a distinct advantage when the RNA is embedded in a larger macromolecular context such as RNA-protein or RNA-ligand complexes. ^20^ By explicitly accounting for the surrounding environment, AF3 achieves modest yet consistent gains in recapitulating key RNA contacts, particularly non-Watson-Crick (NWC) base pairs^20^. In fact, AF3 surpasses most tools by a substantial margin for these NWC interactions^21^, although the absolute performance remains modest, indicating that further methodological advances are required to model RNA-specific contacts reliably. However, on a broad set of challenging benchmarks, AF3 is outperformed by the specialized framework DeepFoldRNA^21,22^ and by RhoFold+,^23^ underscoring the ongoing need for improvement across diverse RNA prediction scenarios. That said, AF3 is the sole method identified among those benchmarked (including deep learning, ab initio, and template-based approaches) that can successfully process long sequences (from 200 up to 3000 nucleotides). It performs well for specific targets as described in the DR2Fold benchmark paper^24^.

### I.B. Deep-learning frameworks for RNA structure prediction in macromolecular complexes

Several deep-learning-based frameworks can now predict RNA structures in the context of macromolecular assemblies, i.e., they model the RNA chain together with its surrounding partners (proteins, DNA, other RNAs, or small-molecule ligands)^21^:

- **AlphaFold-3 (AF3):** Explicitly engineered to handle both isolated RNA chains and RNA–protein complexes, AF3 has been extended to accommodate heterogeneous assemblies that may contain proteins, DNA, ligands, and additional RNA strands. Its performance on complex-centric benchmarks has been reported in recent comparative studies^21^.
- **RoseTTAFold-NA (RF2NA):** An adaptation of the RoseTTAFold architecture, RF2NA is capable of modelling not only solitary RNAs but also multi-component systems comprising arbitrary combinations of proteins, DNA, and RNA. Systematic evaluations have demonstrated its competence in predicting RNA structures within such complexes^17,21^.
- **AF3-inspired successors:** Recent methods that inherit AF3 architectural principles and are expressly designed for complex prediction include **Boltz-1** and **HelixFold-3 (HF3)**^19^. These approaches further broaden the scope of RNA-centric modelling by integrating heterogeneous partners directly into the prediction pipeline.

Collectively, these tools illustrate a growing trend towards unified, context-aware RNA structure predictions, moving beyond isolated-chain modelling towards realistic representations of biologically relevant macromolecular assemblies.

The TrmK–tRNA complex provides a stringent testcase for evaluating the ability of AF3 to model RNA-protein interfaces. Indeed, although high-resolution three-dimensional coordinates are available for both components individually, there is no structure of the complex in the PDB, i.e. it is absent from AF3 training set. In a blind test, we compared the five models predicted by the AF3 webserver against the experimentally-validated model of this complex.^25^ Noteworthy, TrmK is a m1A22 tRNA methyltransferase, thus the A22 in the tRNA D-arm must be placed in the TrmK active site. AF3 failed to recapitulate the native RNA-protein interface. Only two of the five models display a modest overlap with the TrmK region observed in the experimentally-validated model, and none approach its correct orientation with respect to the tRNA and the A22 nucleotide to be methylated. Per-residue confidence (pLDDT) remains relatively high for the isolated protein domain but collapses sharply at the interface, offering little predictive insight for either the RNA or the RNA/protein interface (Fig. 1).

**Figure 1:**
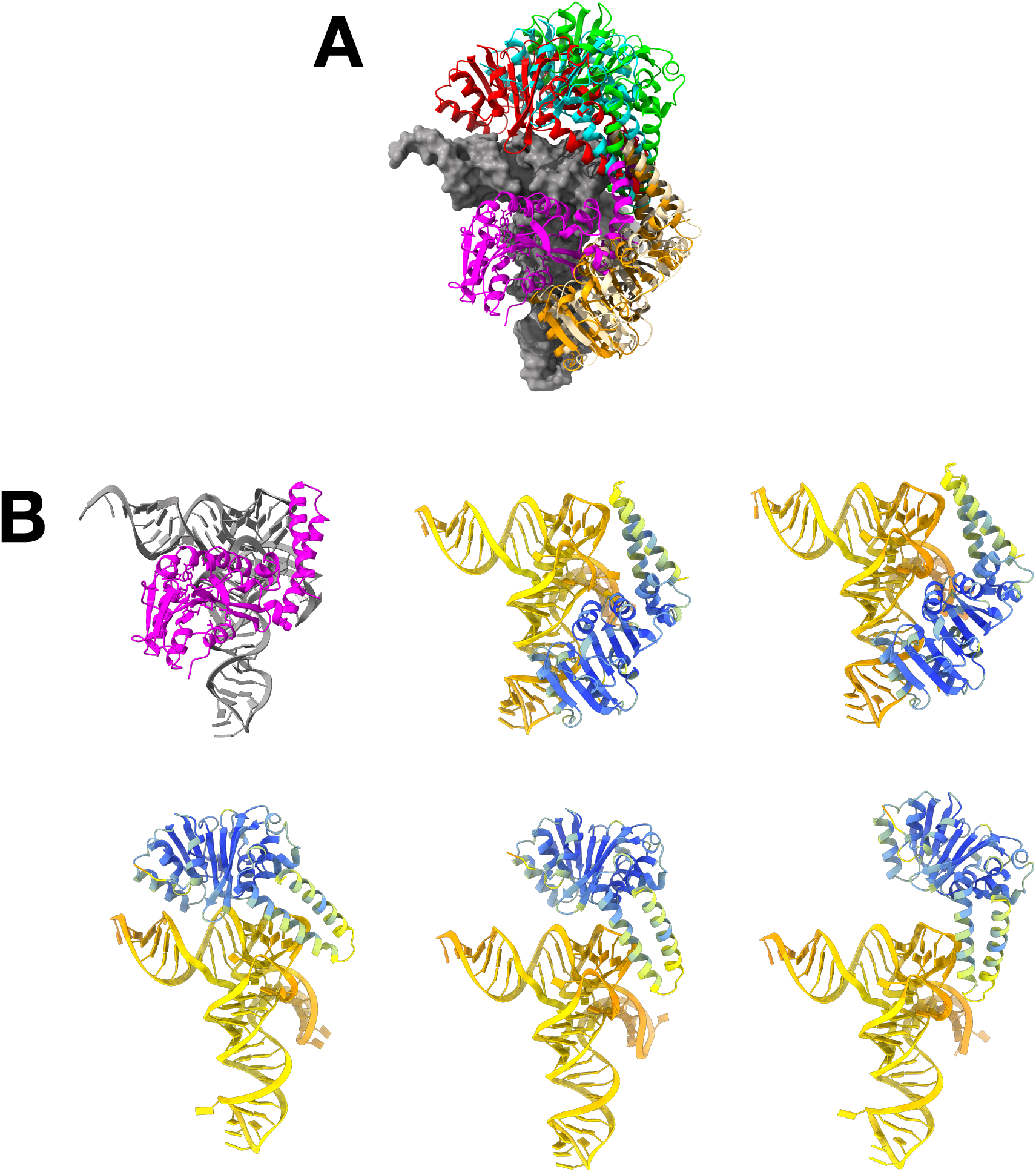
Structural comparison of the experimentally-validated model of the TrmK–tRNA complex with five AF3 predictions. (A) Superposition of the experimentally-validated model (tRNA shown in gray, TrmK in magenta) with the five AF3 models. The models were aligned to the experimental structure using the tRNA (rendered as a solid surface) as the reference for RMS fitting. (B) Side-by-side view of each individual AF3 model (colored according to its per-residue pLDDT confidence score) together with the corresponding experimentally-validated model. This panel highlights local variations in model confidence and structural deviations across the five predictions.

### II. Protein transmembrane domain prediction

Mitofusins are complex membrane proteins with large domains and low-resolution experimental data and therefore constitute a challenging test case for deep-learning prediction. Their transmembrane domain is particularly difficult to model and is poorly predicted by AlphaFold-2 with low pLDDT scores and inconsistent helical geometry.^26^ Improvements can however be made thanks to the new possibilities offered by new models. Providing additional input data such as custom MSAs allows for more reliable predictions,^27^ but the conformational diversity remains limited, as AF only predicts the open conformation of mitofusins, limiting its ability to capture functional dynamics (e.g., closed/tethered states involved in membrane fusion). This underlines a broader issue: current AI tools struggle with modeling large-scale conformational transitions critical to function,^28^ and often require specific tuning such as MSA depth reduction or template-guided predictions in order to obtain conformational diversity.^29^

Multiple C2 Domain and Transmembrane Region Protein (MCTP) proteins are membrane proteins involved in the regulation of plant plasmodesmata, which are intercellular channels allowing communication and molecular transport between cells. Their structure is largely unknown and thus modelling is crucial to their study. Recent work by Sritharan et al. on their transmembrane domains showed that while deep-learning predictors can provide a small variety of conformations, most of the conformational landscape is still not explored by the current agents.^30^

For both mitofusins and MCTPs, while deep-learning predictions can act as a valuable starting point for transmembrane domain determination, the use of molecular dynamics simulations in order to truly explore their conformational landscape was still necessary. ^30^ These examples thus highlight the importance of dynamics for structural predictions, which will constitute one of the next frontiers in biomolecular modelling.

### III. Explicit ion modelling in protein prediction

The possibility to explicitly include ions in AF3 predictions opens new possibilities for modeling biomolecular interactions and conformational changes that depend on ionic environments. This feature can improve the prediction of binding sites and functionally relevant conformations, such as enzyme active states. However, the approach has limitations: ion placement and induced conformational changes may be biased or inaccurate due to gaps in training data, sometimes resulting in low-confidence predictions or non-native states. The impact of the ion type and stoichiometry on model accuracy also requires careful consideration and validation, ideally supported by experimental data or complementary computational methods. Here we assess how accurate explicit ion modelling is, in terms of placement for case of MCTPs and in terms of conformational switch for the adenylate kinase (Adk) protein.

### III.A. MCTP proteins: when the binding site is unsure

Members of the MCTP family are characterized by three to four C2 domains and two transmembrane regions that are structurally conserved across eukaryotes, from invertebrates (Drosophila, C. elegans) to plants (Arabidopsis thaliana) and mammals. ^31–33^ In invertebrates and mammals, these C2 domains interact with calcium. ^31,32^ However, in plants, it remains unclear whether the C2 domains retain this calcium-binding ability. In Arabidopsis thaliana, MCTP proteins are involved in organizing and regulating plasmodesmata, specialized channels facilitating intercellular communication. Since calcium acts as a secondary messenger in plant cells, influencing various signaling pathways,^34^ computational tools such as AlphaFold3 (AF3) can be employed to predict the structures of plant MCTP C2 domains and identify potential calcium-binding sites. However, AF3 predictions may include inaccuracies or “hallucinations”, raising the question of how to reliably confirm these binding sites.^4^ To address this, we propose combining AF3 with a panel of complementary computational tools, such as molecular dynamics simulations, electrostatic protein profile and binding site prediction algorithms related to experimental data, to enhance the accuracy of identifying calcium interaction sites in plant MCTP proteins.

In the top-ranked AF3 model generated with one calcium ion, the predicted local distance difference test (pLDDT) score for calcium placement was low (pLDDT = 43.25 and ipTM = 0.62), indicating low confidence in the predicted calcium-binding site. To explore the impact of calcium stoichiometry, we varied the number of calcium ions from 1 to 10 during AF3 inference (see Figure 2). While the predicted C2B structure remained similar regardless of calcium presence or number (indicating robust structural prediction), increasing the number of calcium ions improved the confidence in calcium-binding site predictions, as reflected by higher pLDDT scores. This trend aligns with the observation that experimental structures in AF3 training database often include multiple calcium ions for proteins with C2 domains. Among RCSB structures annotated as C2 domains, the number of calcium ions per structure is variable, with many structures containing no calcium and others containing one or several calcium ions (see data in the Zenodo archive: https://zenodo.org/records/21534689). Since calcium occupancy is heterogeneous among experimentally resolved C2 domain-containing structures, AlphaFold 3 predictions involving calcium should be interpreted as conditional models depending on the number of ions provided as input, rather than as an unbiased prediction of calcium stoichiometry.

**Figure 2:**
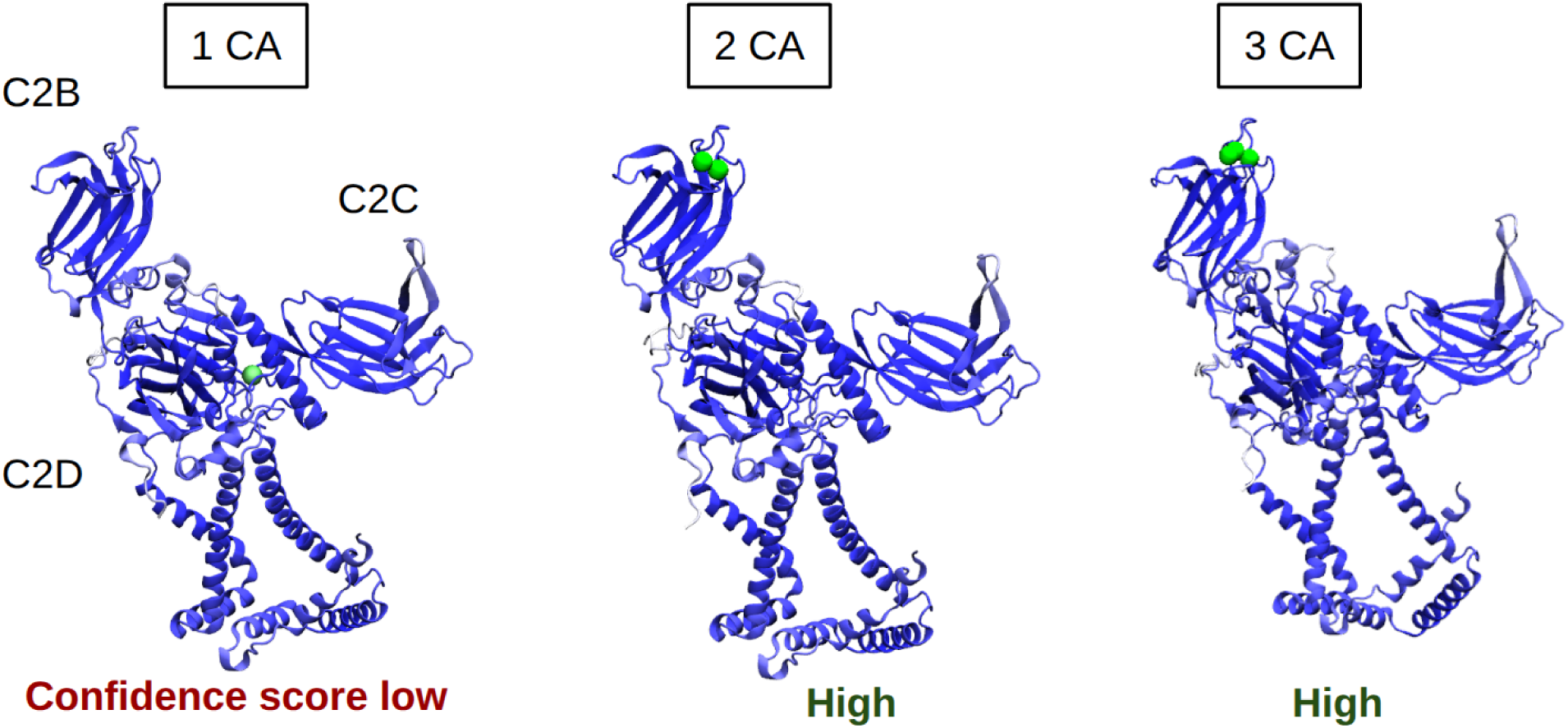
AF3 predictions of the calcium-binding site in MCTP4 (model 0 shown for each condition). MCTP4 is colored in blue, calcium cations in green. Model with one calcium ion: pLDDT = 43.25, ipTM = 0.61. Model with two calcium ions: pLDDT = 60.83, ipTM = 0.86. Model with three calcium ions: pLDDT = 67.73, ipTM = 0.81.

To further assess the uncertainty regarding calcium binding, we employed AlphaFill to examine whether calcium cations could be transferred onto the predicted C2 domain structure based on similarity with experimentally-resolved structures. ^35^ AlphaFill enriches AlphaFold models by transplanting ligands, cofactors and metal ions from homologous experimental structures, hereby providing additional structural context for functional interpretation. In addition, we performed an electrostatic potential analysis using the Adaptive Poisson-Boltzmann Solver (APBS). This analysis revealed an electronegative region in the C2B domain, comprising five aspartate residues, suggestive of a potential calcium-binding site (see Figure 3). These findings provide preliminary evidence of a calcium-binding region in MCTP4 C2B domain.

**Figure 3:**
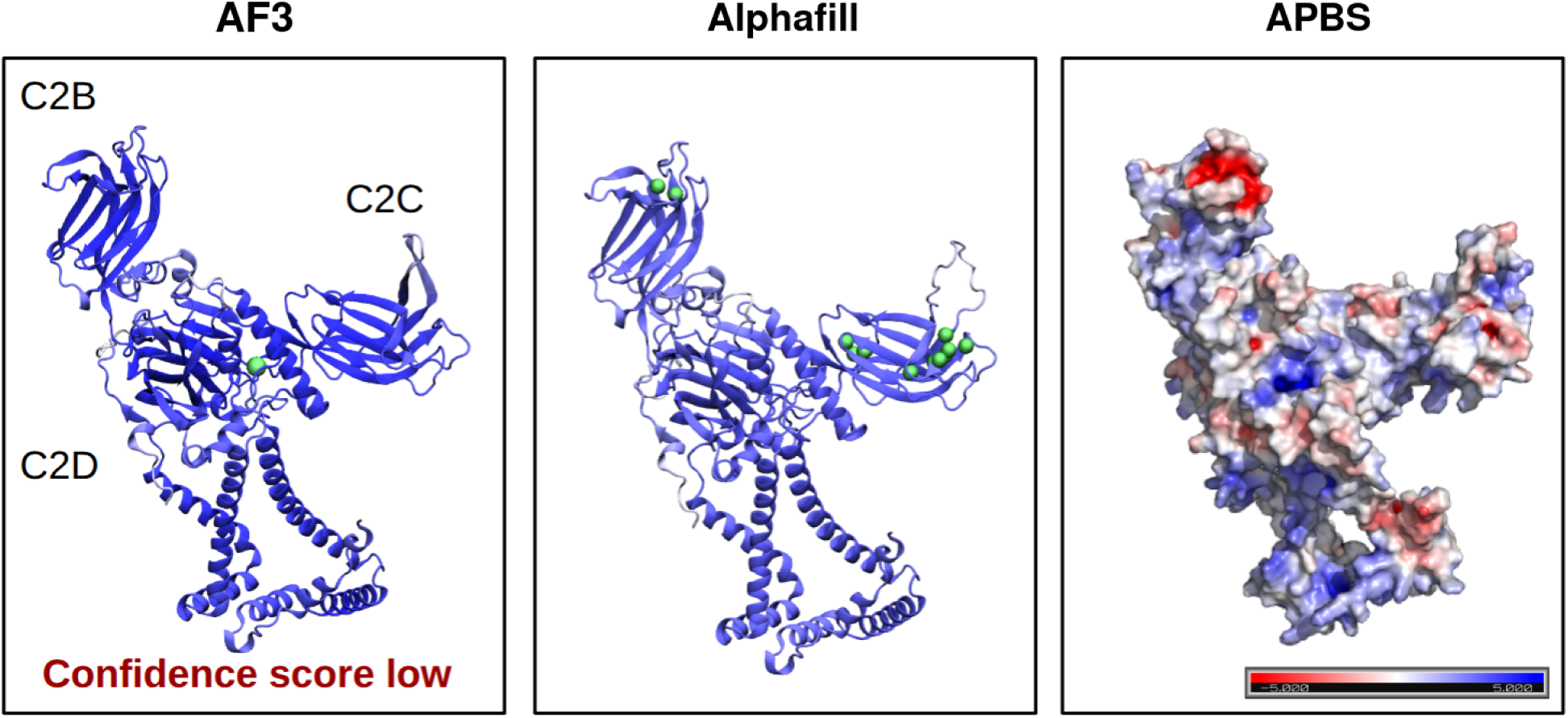
Various methods to predict the binding site of calcium on the MCTP monomer, from purely deep-learning-based on the left to purely physics-based on the right.

### III.B. Conformational changes associated with ion binding in adenylate kinases

While recent deep-learning tools such as AF2, RosettaFold, and OmegaFold have demonstrated remarkable accuracy in predicting ordered protein structures, their ability to capture the dynamic conformational landscape of protein switchers is still limited. ^36,37^ AlphaFold tends to return a single conformational state, and alternative conformations can be obtained by perturbing the model input or sampling.^38,39^ These methods predate AF3 and act on the model rather than on the physical system; here we perturb the physical input instead. We report a strategic example using AF3 showing that the inclusion of ions during model prediction (a new feature of AF3) can induce a distinct conformation in the well-studied adenylate kinase (Adk) protein. Adk is a key enzyme involved in maintaining cellular energy balance by reversibly transferring phosphate groups between ATP, ADP, and AMP undergoing conformational transitions between open and closed states. ^40^ We predicted its structure with AF3 in four different scenarios; without ions or ligand, with ions, with ligand, with ions and ligand. For the structure predictions, different combinations of ions (Zn^2+^, Mg^2+^, K^+^, Cl*^−^*), and ligands (ATP, ADP), were explored. The specific ion and ligand composition varied between organisms and prediction trials, with each model containing between 0 and 4 ions and 0 and 1 ligand.

Overall, we observed that the explicit addition of ions shifted the prediction toward an alternative state (Figure 4). AF3 always predicted an open conformation when only ions were included, while a closed conformation was sampled in the three other scenarios. Figure 4 summarizes these findings across Adk homologs from various organisms, all showing consistent ion-dependent structural differences. The left panel depicts the distance between two selected representative residues from the NMP and the LID arm of Adk for different homologous protein models of AF3. Residue pairs were selected after structural alignment of the Adk proteins to compare equivalent positions across organisms (where the selected pairs are Gly42–Leu157 for *B. stearothermophilus* and *G. psychrophilus*, Glu42–Leu157 for *B. subtilis*, Gly42–Val147 for *A. aeolicus* and Gly42–Leu153 for *E. coli*). A distance above 35 Å is here taken to define an open state and a smaller distance a closed one. This cutoff is a practical threshold chosen to separate the two populations based on the state separation observed between experimental structures (Figure 4, right pannel), rather than a standard order parameter, as the open/closed transition is more commonly described by the LID–CORE and NMP–CORE inter-domain distances. ^41,42^ To further understand these predictions, experimental models available for the investigated organisms were also plotted as a reference on the right panel applying the same residue selection. Contrary to what is predicted by AF3, it is clear that in the experimental data an open conformation exists in different cases (with only ions, with only ligands, without ions or ligands). This observation may indicate a tendency of AF3 to favor specific types of conformations in the presence of ions for Adk and potentially, similar fold-switching proteins.

**Figure 4:**
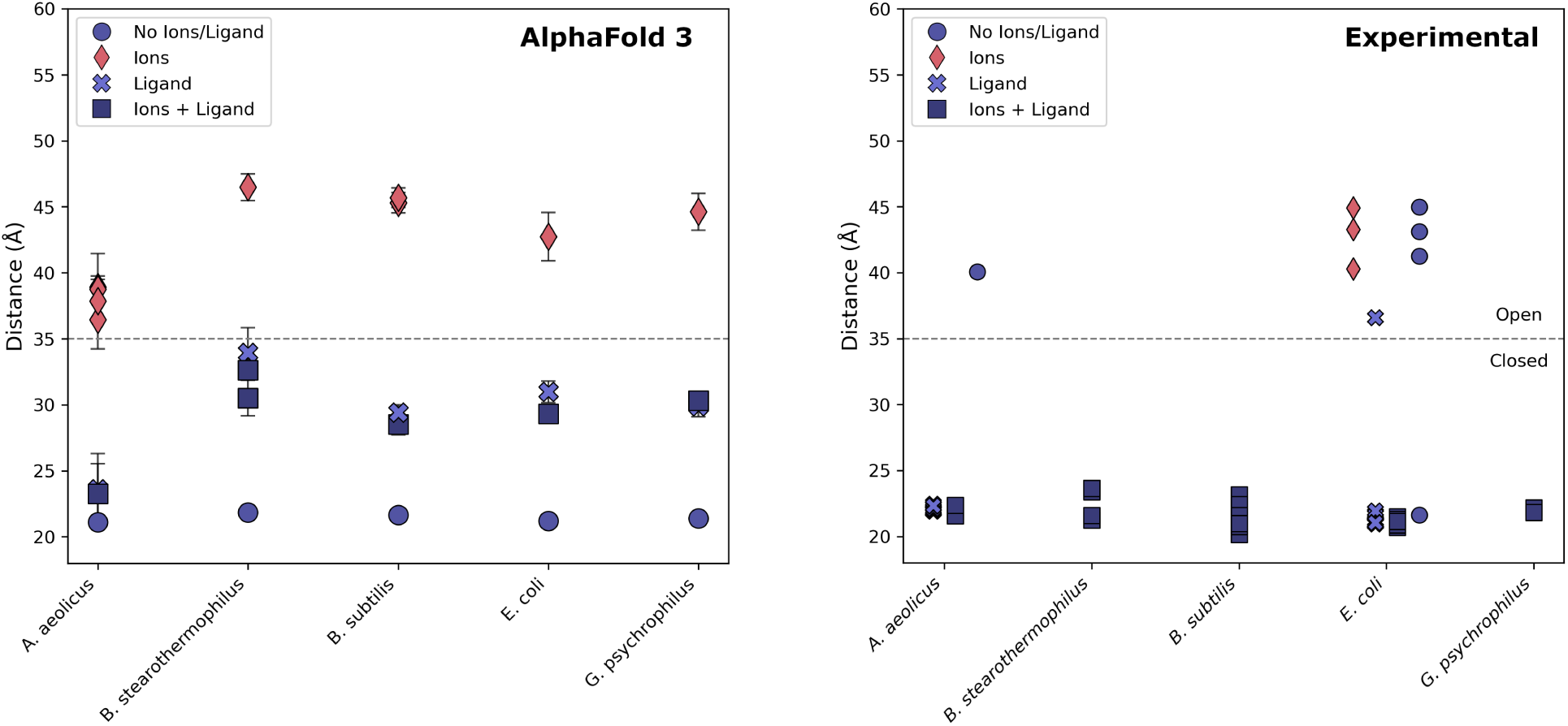
Distance between two key residues of the LID and NMP of Adk proteins of A. aeolicus, B. stearothermophilus, B. subtilis, E.coli, G.psychrophilus. Key residues are Gly42–Leu157 for *B. stearothermophilus* and *G. psychrophilus*, Glu42–Leu157 for *B. subtilis*, Gly42–Val147 for *A. aeolicus*, Gly42–Leu153 for *E. coli*. Left pannel: AF3 predictions with four possible initial submissions, (i) without ions or a ligand, (ii) with ions, (iii) with ligand, (iv) with ions and ligand. Error bars are determined from the 5 predicted models by AF3. Right pannel: Experimentally available PDB structures (see Table 1) categorized according to the presence/absence of ions and ligands. The dashed line at 35 Å seperates open and closed conformations of the PDB structures, and is chosen as an indicative threshold.

**Table 1:**
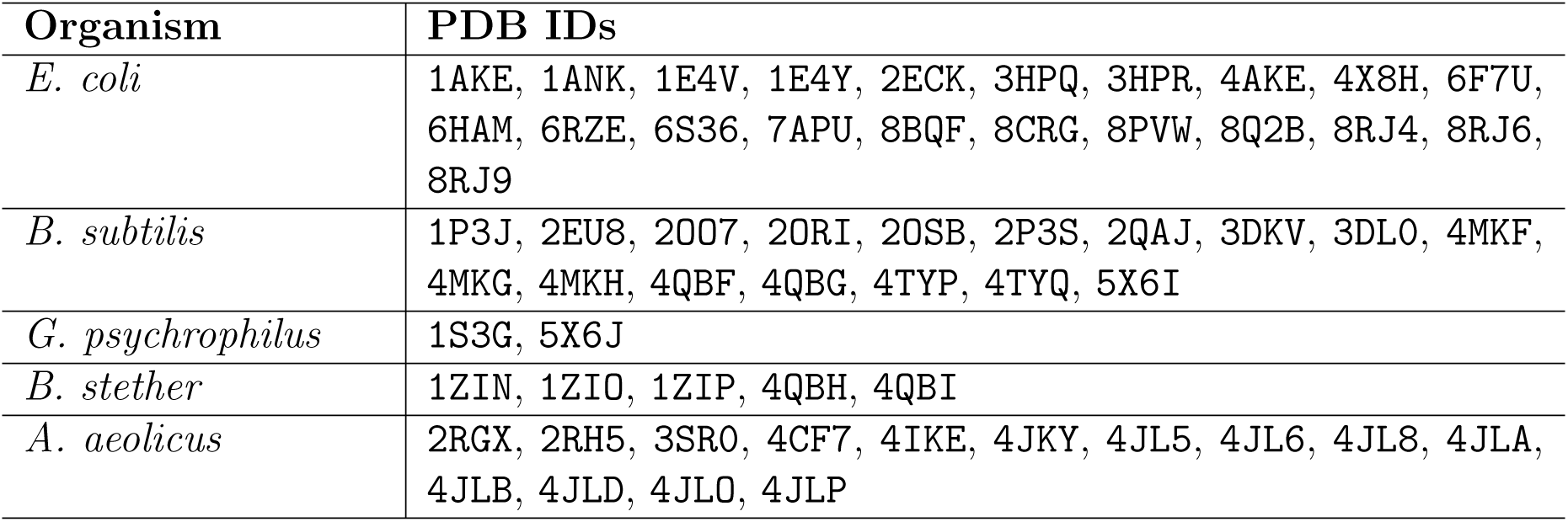
PDB structures of the ADK protein homologs available for each organism.

From these examples, we demonstrate that it is possible to trigger the sampling of distinct conformational states by addition of ligands and in particular ions to the prediction query. The placement of these ions however remains highly unreliable, and further considerations on this topic will be explored in the next part of this perspective.

### Persistent modeling limitations and how to circumvent them with explicit ion addition

Deep-learning tools still present clear limitations in their ability to faithfully model biomolecular systems outside of the common folded protein. We here discuss such limits in particular for nucleic acids and protein disorder, but also present a novel capability of explicit ion addition to shift the predictions in various ways.

### I. RNA 3D structure prediction is hampered by data-driven limitations

RNA structure prediction is still a major challenge for biomolecular predictors, which particularly struggle to generalize past their training set^18^. These shortcomings were highlighted repeatedly in the RNA-Puzzles and CASP assessments^17,43^. For instance, the fragment-assembly pipeline FARFAR2, which consistently ranks among the top non-ML predictors, struggles with RNAs longer than ∼80 nucleotides because sampling becomes insufficient and the scoring function frequently misranks near-native conformations^44^. Recent work has shown that a DL-derived scoring term (ARES) can better predict the root-mean-square deviation (RMSD) to the unknown native state^45^, yet the underlying data gaps remain a major obstacle. We provide a detailed description of the data-driven limitations associated with RNA structure prediction in Table 2.

**Table 2:** Key data-related limitations that affect RNA 3D structure prediction.

| Issue | Details | Impact on Modelling |
| --- | --- | --- |
| Sparse high-resolution RNA structures | $\sim 25\times$ fewer total entries and $\sim 110\times$ fewer structures resolved at $<2.0\text{\AA}$ compared with proteins<sup>46</sup>. Flexible RNAs also exhibit higher B-factors, reflecting poorer local resolution. | Reduces statistical power of template-based and DL models; limits learning of fine-grained geometric constraints. |
| Incomplete or biased sequence alignments | RNA families are under-represented in Rfam; consensus secondary structures are often poorly folded, providing weak constraints for 3D reconstruction. | Degrades MSA-driven predictors (e.g., RoseTTAFold-RNA) and hampers extraction of co-evolutionary signals. |
| Structural complexity | non-Watson-Crick pairs are highly constrained in space | local error leads to cumulative errors in neighboring residues |
| Backbone flexibility and conformational sampling | backbone has seven torsion angles | many acceptable alternative conformations, but these alternatives place non-helical nucleotides incorrectly |
| Insufficient enforcement of standard stereochemistry | chirality violations | incorrect local fold or backbone orientation |
| Reliability of energy-based scoring functions | mis-ranking of decoy models versus near-native structures | scoring methods fail to consistently selecting the best structure among model predictions |
| Multimeric assembly and stoichiometry as an added challenge | computational overload, memory failure / dynamic conformational changes | lower prediction accuracy and confidence |

Despite encouraging trends, DL models have not yet delivered uniformly high-accuracy predictions across the full spectrum of RNA fold families. Key obstacles include:

- **Training-set bias** — Structures that are closely represented in the training corpus receive markedly higher TM-scores.
- **Scarcity of high-quality 3D data** — The limited number of experimentally resolved RNAs constrains the diversity of learned representations.
- Role of metal ions, solvent effects, and post-transcriptional modifications

— These factors may be important for obtaining geometrically accurate RNA models.

Addressing these challenges will require coordinated initiatives to (i) enlarge and curate high-quality RNA-structure repositories, (ii) deepen our understanding of the chemical environment surrounding RNA such as including metal ions, solvation shells, and post-transcriptional modifications, (iii) integrate high-throughput experimental data on primary sequences and secondary-structure ensembles, and (iv) develop hybrid pipelines that judiciously blend machine-learning components with physics-based or template-based (non-ML) sampling and scoring modules, choosing the optimal strategy according to the similarity between the target RNA and the families represented in the training data^47,48^.

The transition from classical fragment-assembly and physics-based protocols to ML-driven RNA structure prediction mirrors the trajectory observed for proteins. Yet, the scarcity of high-resolution RNA structures and the uneven coverage of RNA families impose a ceiling on current model performance. Although ML methods—especially those that exploit contextual information—already outperform many non-ML approaches, their advantage diminishes or vanishes for novel or orphan RNAs. Addressing the initiatives outlined above will be necessary before generative AI can achieve, for RNA, the same predictive reliability it has reached for proteins.

### II. Explicit addition of ions to circumvent limits in predictions

#### II.A. AF3 responds to ionic addition in nucleic acid predictions

In order to go further than bare sequence modeling and to assess whether nucleic acid predictions with explicit ions are accurately modelled by AF3, we conducted an explorative work on the structures of 2 G-quadruplexes sequences, inspired by Ochoa and Milam who showed in a groundbreaking study that incorporating explicit ionic cofactors in AF3 modelling could drastically modify the predictions of G-quadruplexes.^15^ A G-quadruplex is a helical structure formed by a nucleic acid sequence, usually stabilized by one or several cations interacting with the bases.

Using AF3, we started by modelling the sequence of the telomeric G-quadruplex 5’-AGGGTTAGGGTTAGGGTTAGGG-3 corresponding to the 1KF1 PDB structure, not studied by Ochoa and Milam. This G-quadruplex is stabilized by three potassium cations, and is one of the first quadruplexes resolved at high resolution. We thus expected this well-known structure to be easily reproduced by AF3 as it was part of its training set. Figure 5 displays the RMSDs of the C1’ atoms between the best prediction of AF3 and the reference 1KF1 PDB structure. Without explicit ion addition, the prediction is very remote from a G-quadruplex conformation. The addition of a single cation is not enough to trigger the conformational shift, and chloride addition never yields a G-quadruplex conformation, indicating that AF3 does capture ion specificity to an extent. Modelling the nucleic sequence with the three expected potassium does create the expected structure, however we did notice that AF3 placed two of the potassium at the same location instead of aligning them (see illustration on the bottom right of Figure 5). This extreme steric clash in the prediction is concerning, we thus wondered what would happen for a sequence that AF3 has not seen before.

**Figure 5:**
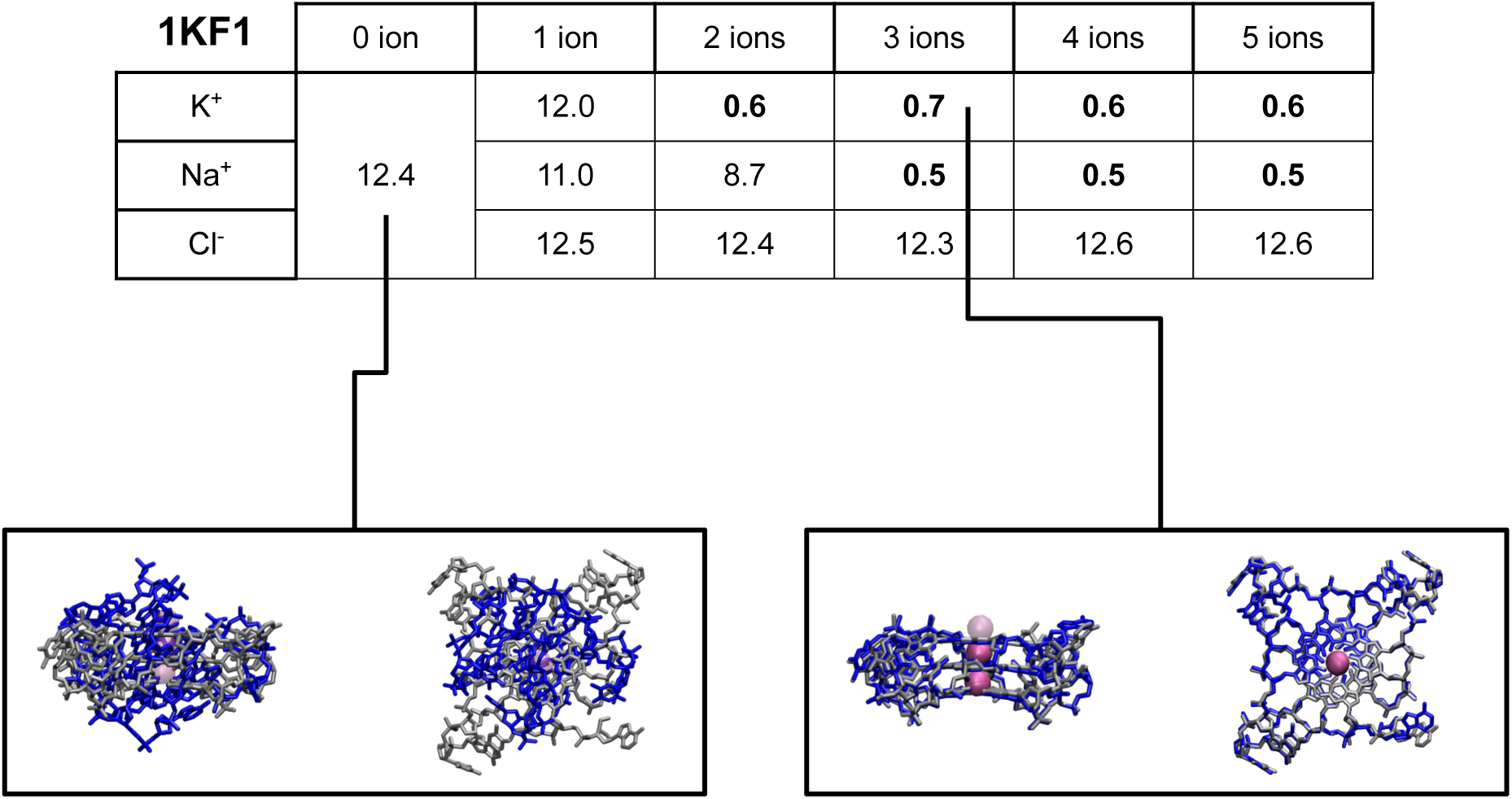
Table of the computed RMSDs of the C1’ atoms in angstroms between the best model predicted by AF3 and the 1KF1 PDB structure of the G-quadruplex 5’-AGGGTTAGGGTTAGGGTTAGGG-3. RMSDs below 2 angstroms are in bold. The superposed structures constitute relevant examples of predictions with respect to the reference. The predicted structure by AF3 is in blue, the reference 1KF1 is in silver. Potassium cations are represented by Van der Waals mauve spheres, opaque for the prediction and transparent for the reference.

Late 2025, Geng et al. successfully crystallized the G-quadruplex sequence 5’-GGGGCCGGGGCCGGGGCCGGGGCC-3 and showed that it could take a dimeric parallel conformation (PDB 9UK6) or a monomeric antiparallel conformation (PDB 9UK8). ^49^ Given that these crystal structures were published later than the release of AF3, we can safely assume that they were not present in the training data of the AI model. We thus computed the RMSD of the C1’ between the best monomeric prediction by AF3 and the PDB reference structures (see Figure 6). None of the predicted conformations adopted an antiparallel structure as all computed RMSDs with PDB 9UK8 were above 7.5Å (data not shown). While the prediction without explicit ions did not match the parallel structure either, addition of ions did allow for the generation of closer conformations to the parallel reference PDB 9UK6. However, good agreement with the reference was only achieved through addition of two to four chloride anions, in contradiction with the stabilizing potassium cations present in the crystal. Furthermore, these anions were not placed in lieu of the cations, but in a symmetric pattern at the periphery of the structure. This bias towards symmetry in ion positioning was observed in multiple predictions, and might constitute an interesting feature learned by AF3 during its training.

**Figure 6:**
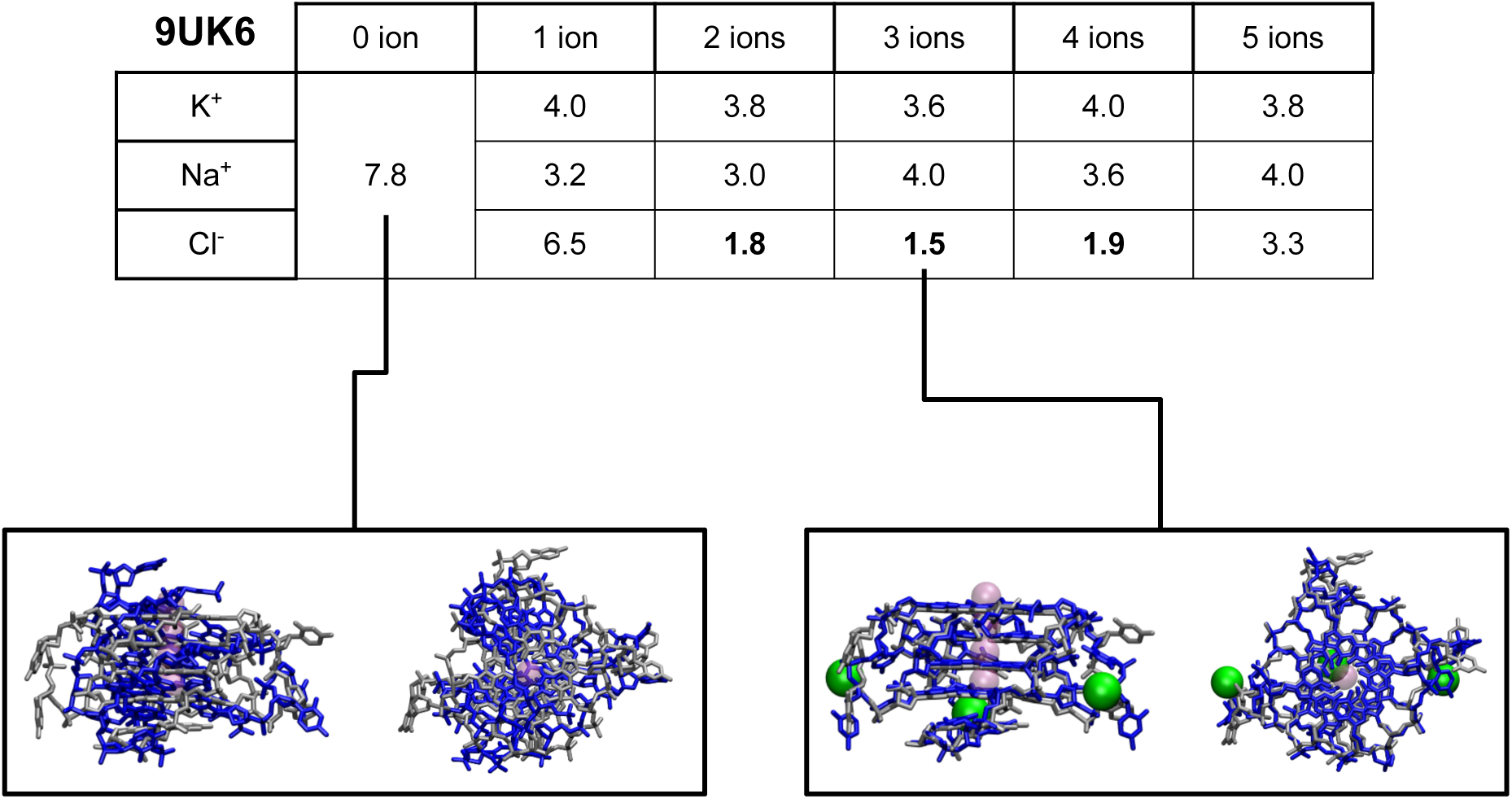
Table of the computed RMSDs of the C1’ atoms in angstroms between the best model predicted by AF3 and the 9UK6 PDB structure of the G-quadruplex 5’-GGGGCCGGGGCCGGGGCCGGGGCC-3. RMSDs below 2 angstroms are in bold. The superposed structures constitute relevant examples of predictions with respect to the reference. The predicted structure by AF3 is in blue, the reference 9UK6 is in silver. Potassium cations are represented by Van der Waals transparent mauve spheres for the reference, and chloride anions by opaque green spheres for the prediction.

Overall, this attempt at G-quadruplex predictions with AF3 yielded mixed results even for easy targets, and showed that further refinements are needed in order to accurately capture the biochemistry of ion-nucleic acid complexes.

#### II.B Calcium-activated allostery still evades AF3 in the absence of known homologs

Human-expressed EndoU (hEndoU) is a RNase known to cleave single-stranded RNAs with a poly(U)-sequence specificity.^50^ hEndoU is activated by a calcium-dependent mechanism, during which the enzyme has been crystallographically shown to bind to four calcium ions (PDB structure 9FTW).^51^ The activation process of hEndoU is believed to be mediated by an allosteric mechanism,^52^ however no experimental structure of the inactivated form is available to this day. Additionally, while the cristallographic structure 9FTW displays five putative calcium binding sites, the fifth site has been shown to be an artefact of the cristallization process. We thus tried to trigger the conformational change from inactive to active form of hEndoU by modelling its sequence with AF3 in the absence and presence of 1 to 5 calcium cations. Figure 7 reveals that despite the removal of calcium cations, AF3 was not able to produce an inactive form of the enzyme. However, AF3 correctly fills up the putative binding sites by starting from site 3, then site 1, then site 2, and finally site 4 as the number of calcium ions is increased. Interestingly, when adding a fifth cation, AF3 does not populate the fifth binding site, but instead models it between sites 1 and 2. These observations lead us to believe that the explicit addition (or lack thereof) of ions alone is not sufficient to trigger a conformational change if AF3 does not have knowledge of a homolog, as was the case for Adk proteins. However, AF3 seems to correctly identify the correct binding sites of the hEndoU enzyme, despite the crystallization artefact. These results are encouraging for the use of AF3 to detect ligand binding sites in allosteric proteins, but progress remains to be made regarding the prediction of active and inactive states without available homologs.

**Figure 7:**
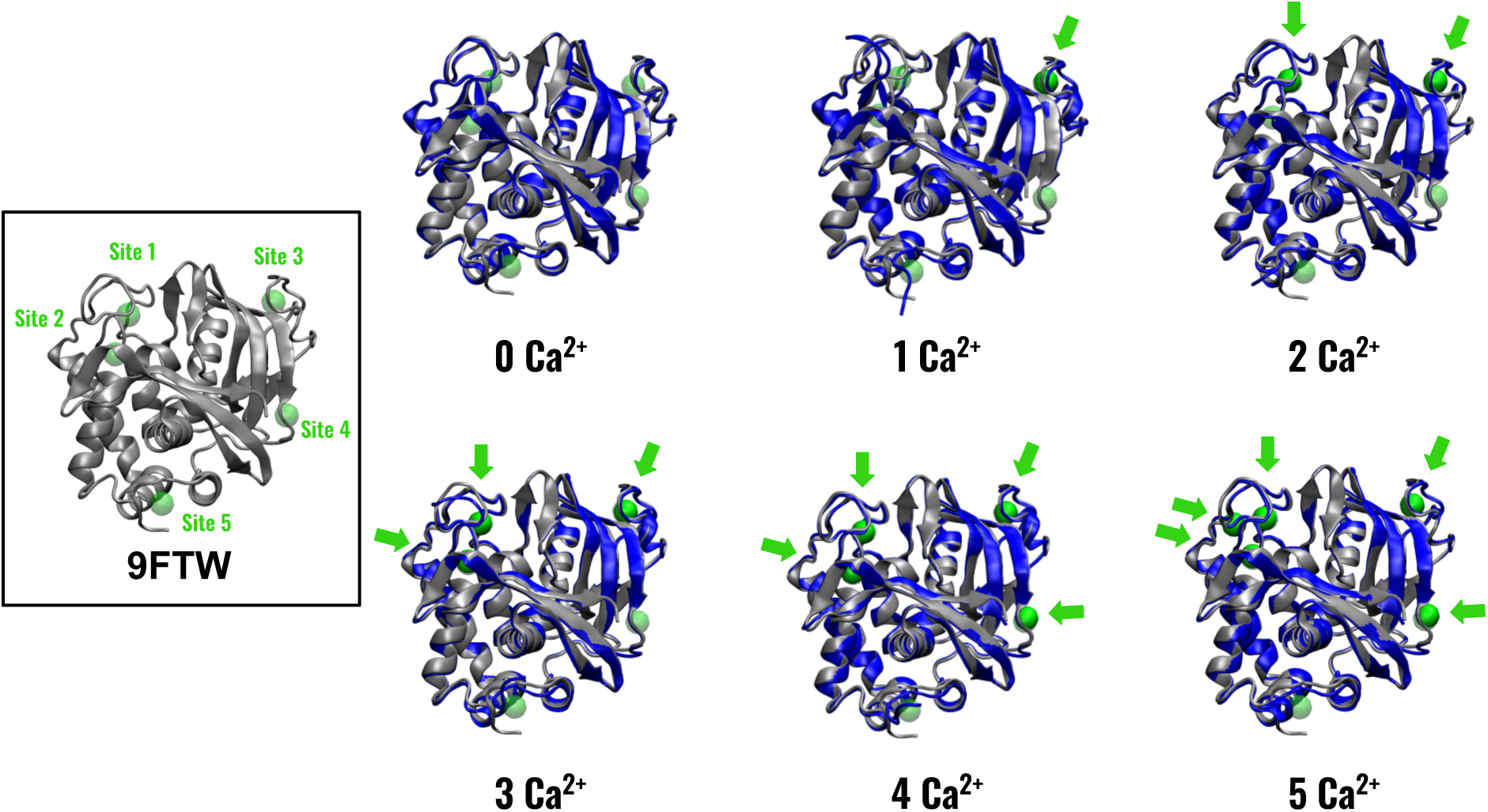
AF3 predictions of the structure of enzyme hEndoU in absence and presence of calcium cations. Calcium binding sites are indicated in green in the box containing the crystallographic structure on the left. Green arrows point to where the calcium cations are predicted by AF3 to be binded. The predicted protein structure by AF3 is in blue, the reference 9FTW is in silver. Calcium cations are represented by Van der Waals transparent green spheres for the reference, and by opaque green spheres for the prediction. From 0 to 5 Ca^2+^, RMSDs of the C*_α_*s of the predicted protein with respect to those of the reference are 1.3Å, 1.2Å, 1.1Å, 1.4Å, 1.0Å and 1.0Å.

#### II.C. AF3 folds IDRs and IDPs in an ion-dependent manner

AlphaFold models struggle to accurately fold peptides. ^53^ Their tendency to fold some IDPs and IDRs is also a known concern for the capacity of AF2 and AF3 to correctly predict the folding propensity of proteins.^54^ This limitation is particularly salient for AF3, whose diffusion-based generative architecture introduces a distinct failure mode compared to AF2: rather than defaulting to extended, ribbon-like conformations for disordered regions, AF3 tends to hallucinate spurious local structural order, even though these regions are typically flagged with very low confidence scores. To mitigate this “spaghetti” appearance and recover AF2-like ribbon predictions for disordered segments, the AF3 developers introduced cross-distillation training from AF2 predictions, combined with a ranking term that favors models with greater solvent-accessible surface area. ^4^ Recent works by Bret et al. and Fayetorbay et al. also pinpointed the necessity to carefully treat disorder in proteins when using the AF2 software, highlighting the difficulty for deep-learning predictors to accurately account for disorder in proteins and complexes. ^55,56^ We report here two examples of such limitations, and our findings that the addition of ions within the framework of AF3 allows to modify the prediction without changing the confidence of the model.

**Example 1: The R2 repeat domain of the Tau protein** The Tau protein is a well-studied Microtubule-Associated Protein (MAP) IDP of 441 residues in its longest isoform.^57^ Its role as a stabilizer of microtubules (MTs) in neuronal cells is mostly mediated by its repeat domains. The structure of the tau repeat domain bound to microtubules was recently resolved by cryo-EM, showing that each repeat including R2 adopts an *extended, secondarystructure-free* conformation along the protofilament, spanning three tubulin monomers.^58^ The free protein remains intrinsically disordered and the inter-repeat linkers stay flexible, so R2 is not expected to fold into any regular secondary structure. We therefore expected a peptide with the sequence of the R2 repeat domain to be predicted as disordered by AF3, but to our surprise the model confidently predicts complete folding as an alpha-helix (Figure 8A). Furthermore, the average pLDDT score exceeded 80% on all 5 models presented by AF3 (82.3, 84.4, 84.8, 81.8, 81.7). This prompted us to use the latest feature of AF3, i.e. the explicit inclusion of ions, to study how this new AF version would be sensitive to it. We restrained ourselves to monovalent ions, sodium (Na^+^), potassium (K^+^) and chloride (Cl*^−^*). We will call predictions without ions “dry” predictions in the following discussion, and will use them as a reference. To study the structural changes in the prediction, we use the radius of gyration R*_g_* of the peptides, assuming that the R*_g_* of an alpha-helical prediction would not vary considerably as long as the predicted peptide is folded. This was confirmed by calculating the spread of R*_g_* from the five models predicted without ions (11.9 +/-0.3 Å). We then predicted the structure of R2 with 1,2,3,4 and 5 ions (Figure 8A). Addition of ions resulted in a wider spread of radii of gyrations between the models than for the dry prediction, in all cases but for the addition of 4 chloride anions or for more than 3 sodium cations. The average value also increases compared to the dry prediction in all cases. Addition of K+ causes the strongest unraveling, with the average value of R*_g_* increasing by 50% for 5 K+, and the broadest recorded spread at 2.9 Å.

**Figure 8:**
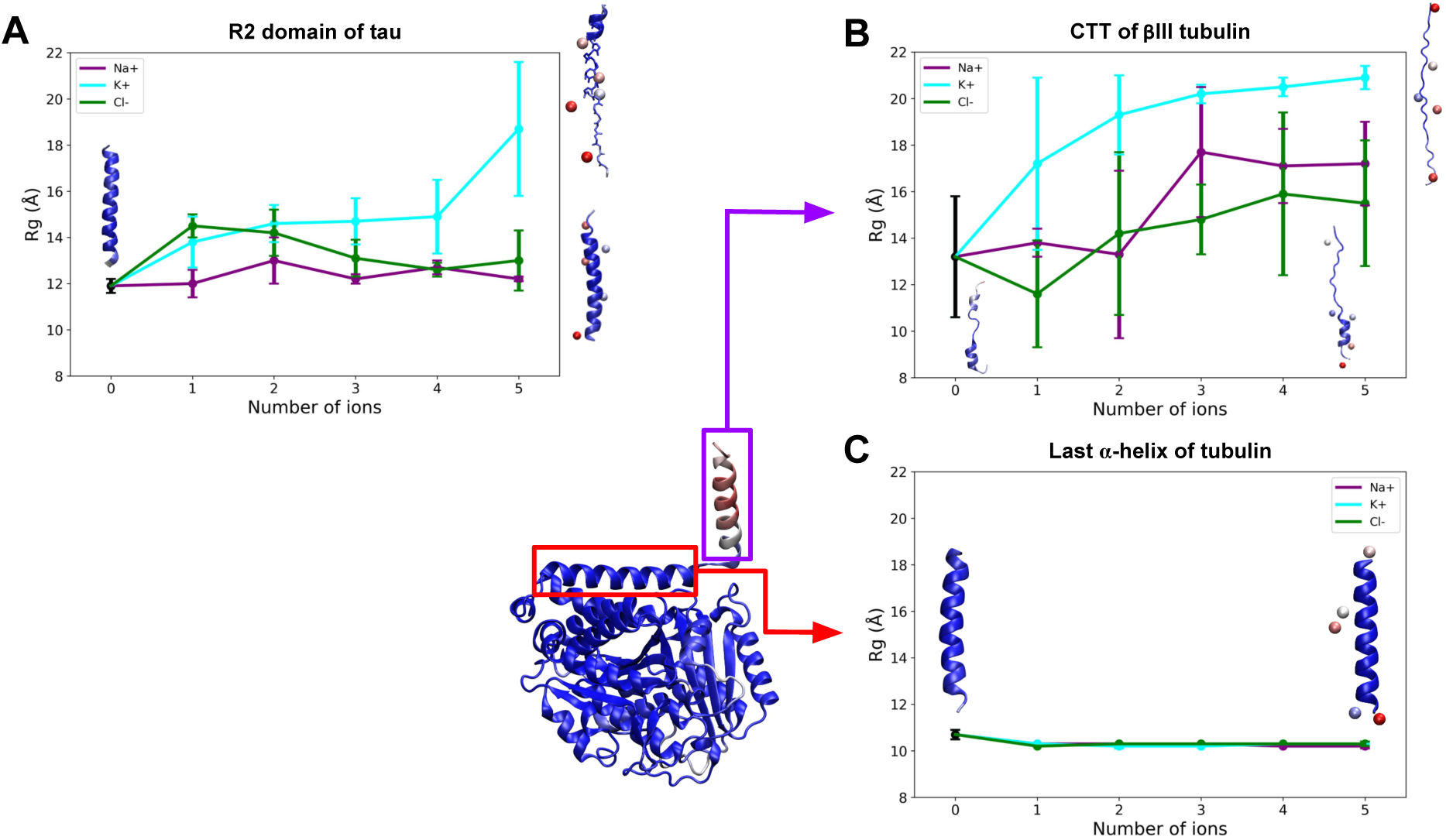
Effect of explicit ion addition on the distribution of radius of gyration R*_g_* for A) the R2 domain of the tau protein, B) the CTT of the βIII tubulin and C) the last α-helix of the same tubulin. The represented tubulin monomer and the illustrative peptide predictions are colored by pLDDT score, where deep red is 0, white is 50 and deep blue is 100.

AF3 thus generates over-folded structures for this IDP, but explicit ionic addition (in particular of potassium) in the input of the deep-learning framework seems to be a practical way to induce the unravelling of the misfolded alpha-helix.

**Example 2: The C-terminal tail of the** β**III-tubulin** We then tested whether this effect would also happen for a misfolded IDR. Tubulins are the constituent proteins of MTs and possess a folded core of around 420 amino acid and a disordered C-terminal tail (CTT) with a length depending on the considered tubulin isotype.^59^ βIII-tubulin is mostly expressed in neurons and its CTT is 24-amino-acid long, with a high proportion of negatively charged residues. The disordered character of the CTT is an experimental and computational fact that contributes to its assumed regulatory function of MT-MAP interactions. ^60,61^ Although the pLDDT never exceeds 60 for the best model on the CTT residues, AF3 still predicts an alpha-helical structure for this CTT, as was recently the case for the prediction of an αI/βI tubulin dimer by Shred et al. (Figure 1A of Ref^62^). We therefore applied the same explicit ionic addition scheme in AF3 as for the tau-R2 peptide to the sequence of the CTT (Figure 8B).

Even the dry prediction reveals an heterogeneity between the models, with a standard deviation of R*_g_* of 2.6 Å. The dispersion of R*_g_* is wider than 1 Å in all cases but cases with 3, 4 and 5 K^+^, for which the peptide is fully unfolded and extended. K^+^ thus seems to be the most disruptive ion for the structuration in AF3. To establish whether these effects are specific to disordered fragments like the CTTs, we also conducted the same predictive study on the sequence corresponding to the last α-helix of the βIII tubulin monomer (Figure 8C).

Addition of all ion types led to a small compaction of the R*_g_* values, and almost no deviation was measured between the models, showing that AF3 was fully confident in the helicity of the sequence regardless of the ion context.

These observations lead us to hypothesize that AF3 might be able to decipher between a stable alpha-helix and a misfolded IDR by exposing a peptidic sequence to an addition of potassium cations.

#### II.D. Discussion on the effects of explicit ion addition in AF3

Deep-learning models such as AlphaFold are more and more known to struggle to generalize their predictive power to complexes involving non-proteic ligands^63,64^ and with the modelling of electrostatic interactions.^65^ In their recent work, Sklar et al. also highlighted the necessity to tune AF3 modelling in a system-specific manner when predicting peptide self-assembly.^66^ We have shown in this work additional examples of AF3 limitations, but also uncovered new possibilities for the use of explicit ionic addition. For both nucleic acid and protein sequences, adding ions and in particular potassium to the prediction usually led to alternative molecular conformations often closer to the biochemical reference. We can think of this process of ion addition as a way to add noise to the prediction. This idea of noise addition to the AlphaFold algorithm to increase conformational variety has already been successfully implemented, and is now routinely done with AF2. In AF3 however, we suggest that one does not need to alter the weights, but simply to add ions to the prediction input to artificially add noise at the diffusion step. This explanation remains at the stage of the hypothesis and further work is needed to confirm and statistically quantify it, but the different examples discussed in this perspective seem to concord towards this view.

**Practically speaking, for any AF3 prediction, we recommend modelling sequences both dry and in the presence of different amounts of potassium cations, as this might help sample more diverse conformations or even uncover misfolded disordered parts.**

## Model assessment as a cardinal issue for deep-learning methods

### I. The key question: Can I trust this model ?

A sound question to ask after considering how deceptive deep-learning predictions can be is how to assess the reliability of a given model. Efforts made in accessibility of deeplearning models such as the AlphaFold3 webserver means that people remote from structural biochemistry are now capable of generating models. Contrary to homology modelling, which requires minimal knowledge, and for which one gets an idea of the model quality thanks to the alignment,^67,68^ it not necessarily as easy to assess the quality of a deep-learned model.

In principle, the validation can be made experimentally or with other physics-based modelling methods such as molecular dynamics, but the current trend of model verification tends to favor integrative approaches. In this section, we expose a guideline to validate models produced by deep-learning tools, and detail the metrics and tools available in order to do so.

### II. An experimentalist perspective – on the misuse and need to report confidence metrics

The emergence of AI-driven structure prediction tools such as AlphaFold^69^ and RoseTTAFold^70^ has opened a new era in structural biology. These methods provide remarkably accurate models of proteins and their complexes, changing the way researchers design experiments. On the other hand, they also help interpret, at the molecular level, the data obtained on their favorite system using a broad range of biological techniques, such as low dimensionality experiments (SAXS, FRET, EPR, MS, cross-linking, etc), or simple functional data on variants carrying point mutations.

However, the ease of access to predicted structures could also lead to an increasing misuse of these prediction tools, with possible over-interpretations based on a lack of knowledge related to the confidence metrics, thereby leading to over-enthusiastic conclusions, in particular related to molecular interactions. As these tools become more and more widespread, a critical evaluation of the models reported in publications is definitely needed. The critical evaluation of the models in light of the confidence metrics has been identified as a crucial practice in several early reports evaluating the potentials of AI-based structure prediction tools.^71–73^

Over the last few years, scientists have become increasingly familiar with these tools, and practices to report confidence metrics of their predictions are often followed. This is facilitated by the recent emergence of didactic material for non-specialists. ^74^ However, there remains a number of problematic behaviors, where the confidence scores of the prediction are neither properly considered nor reported in the publications. To improve the overall conduct of the community, uniform practices for model reporting, including local and global confidence metrics should be adopted in publications. As is currently the case for reporting

X-ray, NMR or cryo-EM structures, scientific journals should require authors to adhere to specific guidelines when reporting models built using AI-based structure prediction tools. We believe scientific publications reporting structural models should report relevant confidence metrics, both local and global, with at least pLDDT and PAE graphs, as well as ipTM scores in case of intermolecular interactions (see below for metrics definitions). These practices should not only help authors to better evaluate and analyze their predictions, and thus improve the use of these tools, but also enable a critical evaluation at the peer review stage, gradually instilling good practice in the different groups of the community from the most expert to the most novice.

We thus recommend that publications include the pLDDT and PAE plots, ipTM scores, and, where relevant, actifpTM or ipSAE values (defined below) to provide readers with as much information as possible to assess the accuracy of a model. For RNA or ligand complexes, additional metrics such as the RMSD to experimental data (if available and only for non-fuzzy parts of the molecules) should also be reported given the lower accuracy of deep-learning schemes on such targets.

### III. Overview of available validation tools

The accuracy and reliability of AI-driven structure predictions such as those generated by AlphaFold and related tools depend critically on robust validation metrics. These metrics not only guide the interpretation of predicted models but also help identify potential errors or overinterpretations.^75^ Below, we review the most widely used confidence scores and additional validation tools, along with guidelines for their interpretation and application.

### **III.A.** Core AlphaFold Confidence Metrics

**pLDDT (predicted Local Distance Difference Test)** The pLDDT score provides a per-residue confidence estimate, ranging from 0 to 100. Values above 90 indicate high confidence in the local fold, while scores below 50 suggest disordered or poorly predicted regions. pLDDT is particularly useful for identifying unreliable regions within a model, such as intrinsically disordered segments or flexible loops. It should however be kept in mind (as was notably exemplified in this perspective) that a high pLDDT does not mean a correct answer but rather an answer for which the predictor is confident, a subtle but profound difference. It can be particularly useful to critically consider the context of the prediction (e.g. is the protein a known IDP?) and to check whether the experimental structure of the folded domain was resolved on its own or in the presence of ligands to interpret the pLDDT.

**PAE (Predicted Aligned Error)** The PAE matrix estimates the expected positional error between residue pairs, providing a pairwise confidence map. Low PAE values (dark regions in the matrix on the AlphaFold webserver) indicate high confidence in the relative positioning of residues, while high values suggest uncertainty. PAE is especially valuable for assessing the reliability of interdomain or intersubunit orientations.

**pTM (predicted Template Modeling score)** pTM assesses the overall quality of the predicted structure, with values ranging from 0 to 1. A pTM score above 0.5 generally indicates a reliable global fold. However, pTM can be misleading for small or multimeric structures, where the score may be dominated by a single, well-predicted chain.

**ipTM (interface predicted Template Modeling score)** ipTM evaluates the confidence in the predicted interfaces of multimeric complexes. An ipTM score above 0.8 is considered highly confident, while scores below 0.5 indicate low confidence. ipTM is more informative than pTM for assessing protein-protein interactions, as it focuses specifically on interface accuracy. However, ipTM can be affected by disordered regions or incorrect interface predictions, and its interpretation should be complemented with PAE and pLDDT analysis.

### III.B. Advanced and Complementary Metrics for Multimer Validation

**pDockQ and pDockQ2** pDockQ is a predicted DockQ score, derived from the number of interfacial contacts and the average pLDDT of interface residues.^76^ pDockQ2 improves upon this by incorporating PAE data, making it more reliable for distinguishing correct from incorrect interfaces, especially in heteromeric complexes.

**actifpTM** actifpTM is a refined confidence metric designed to better handle flexible regions in protein complexes.^77^ It provides a more accurate assessment of interactions involving intrinsically disordered or dynamic segments, which are often misrepresented by ipTM alone. actifpTM is particularly useful for peptide-protein or flexible domain interactions.

**LIS (Local Interface Score) and iLIS** The LIS score evaluates the quality of local interfaces. It focuses on low-PAE contacts by applying a PAE cutoff, which allows it to disregard the influence of disordered regions in either the receptor or the ligand on the overall interaction score. When disorder is present in part of the complex under study, LIS therefore offers a clear advantage over the raw ipTM score. It is especially useful for identifying high-confidence interaction patches within larger, less confident interfaces. ^78^ Using the same filtered PAE region, iLIS further restricts the analysis to interacting residues (distance < 8 Å). iLIS combines the LIS with the cLIS (contact-filtered LIS), which allows for a more robust analysis of interfaces. ^79^

**ipSAE** ipSAE is a recent metric that uses the PAE matrix to calculate a pTM-like score specifically for interfaces. ^80^ In most AI-based prediction software, the predicted aligned error distribution is discarded after calculation, with only the resulting PAE values being saved. This makes it impossible to directly recompute chain-chain ipTM or ipTM restricted to a specific region. Roland Dunbrack proposed a workaround that reconstructs an approximate ipTM matrix from the saved PAE values alone, enabling ipTM to be recomputed on specific residues, domains, or chains. The ipSAE score follows the same logic as LIS, focusing on low-error regions and interacting residues. It aims to address limitations of ipTM by focusing on the predicted aligned error of interface residues, providing a more sensitive measure for distinguishing true from false interactions.

### III.C. Tools for Automated Validation and Reporting

Several tools and webservers now provide automated validation reports, integrating multiple metrics and visualizations for ease of interpretation.

**AlphaFold3 Webserver:** https://alphafoldserver.com/ The server not only allows to model proteins, nucleic acids and a wide variety of ligands, it also provides integrated confidence metrics and visualizations for both protein and multimer predictions, including pLDDT, PAE, pTM, and ipTM.

**ColabFold:** https://github.com/sokrypton/ColabFold This online version of AF2 offers user-friendly access to AlphaFold predictions and includes automated generation of confidence plots and reports.^81^

**PICKLUSTER:** https://gitlab.com/topf-lab/pickluster This ChimeraX plugin that clusters and analyzes protein interfaces, providing access to pLDDT, PAE, ipLDDT, iPAE, pTM, ipTM, and model confidence scores in a single interface.^82^

**ipSAE script:** https://github.com/DunbrackLab/IPSAE Open-source scripts are available for calculating ipSAE and actifpTM from AlphaFold output, enabling advanced interface analysis. Dunbrack provides python scripts to compute ipSAE,^80^ otherwise AF-analysis allow to compute pDockQ1/2, LIS and ipSAE scores whenever the PAE matrix is available.^83^ The advantage of ipSAE over actifpTM is that one can compute it from the PAE matrix, as the actifpTM needs to be computed during the calculation (only in latest versions of Colabfold) and cannot be computed based on PAE matrix.

**AlphaBridge:** https://alpha-bridge.eu/ This webserver allows for fast and easy analysis and visualization of AF3 results, focusing on protein complexes and interfaces.^84^

**ModFOLDdock2:** https://www.reading.ac.uk/bioinf/ModFOLDdock/ The server was used in the CASP15 and CASP16 experiments to assess the quality of models.^85^

**AF-analysis:** https://github.com/samuelmurail/af_analysis This python package allows to analyse AlphaFold outputs in an easy and streamlined fashion. ^83^ AF-analysis provides a GUI linking a Mol* 3D visualisation of the model to pLDDT, PAE and score plots, allowing non-specialists to interactively explore their models alongside the corresponding scores. It also enables the computation of LIS, ipSAE, and pDockQ2 scores, and allows clustering of models generated for the same query.

### III.D. Application to a protein-peptide complex benchmark

Beyond the native AlphaFold confidence metrics, several complementary scores have been proposed to better capture the quality of protein-peptide interfaces, yet their performance varies considerably depending on whether correlations are assessed globally or on a pertarget basis. To evaluate this, we recomputed predictions on the 42 protein-peptide complex benchmark of Bret et al.,^55^ using ColabFold^81^ with default MSA generation and both AlphaFold Multimer v2 and v3 weights,^86^ generating 125 models per query for each weight set (250 models per query in total, 10,500 models overall). Similarly to Bret et al., we obtained models of acceptable quality or better for 40 of the 42 complexes when considering the best DockQ score achieved across all generated models, including 38 complexes reaching medium quality or better and 29 reaching high quality. However, when selecting the top-ranked model for each complex based on ipTM or ranking confidence, only 38 of the 42 complexes reached acceptable quality or better, highlighting that AlphaFold’s own ranking does not always surface the best available model.

When correlated with peptide DockQ across this set of predicted models,^87^ global-topology scores such as pTM (Pearson correlation coefficient r = 0.292) and overall pLDDT (r = 0.439) perform poorly, confirming that metrics designed to assess the full-chain fold are inconsistent to judge peptide positioning. In contrast, interface-focused scores show substantially stronger pooled correlations: ipTM (r = 0.858), ranking confidence (r = 0.849), pdockQ2 (r = 0.687), the inter-chain PAE terms (r ≈-0.73 to-0.76), and the more recently introduced LIS (r up to 0.844 for the peptide to receptor direction). These metrics all track DockQ more closely, as do peptide-restricted variants of ipTM such as actifpTM (r = 0.794) or the peptide-receptor ipTM recomputed using Dunbrack method^80^ that we called ipTM_d0_ (r = 0.779). Notably, the peptide-specific pLDDT (pLDDT_peptide_, restricting the standard pLDDT calculation to peptide residues only) stands out as one of the best pooled performers (r = 0.819), suggesting that confidence in the local peptide conformation itself, rather than in the complex as a whole, is a particularly informative signal.

As shown in Figure 9, both pLDDT_peptide_ and LIS_peptide*−*receptor_ display a clear positive trend with DockQ across the pooled set of models, further supporting their relevance as peptide-specific alternatives to global confidence metrics. However, this apparent strength is misleading: when the same correlations are computed per query (i.e., within the set of 250 models generated for each individual complex, which is the practically relevant scenario for model selection), all scores collapse to a much narrower and lower range (r_query_ ≈ 0.39-0.52), with even the best-performing metrics such as peptide-receptor ipTM_d0_ (r_query_ = 0.518) or actifpTM (r_query_ = 0.599) losing much of their discriminative power. Part of this drop can be explained by the fact that, for certain queries, the 250 generated models show very low structural diversity and are almost uniformly of good quality, leaving little variance in DockQ for any score to explain and artificially deflating the per-query correlation despite AlphaFold consistently predicting the complex correctly.

**Figure 9:**
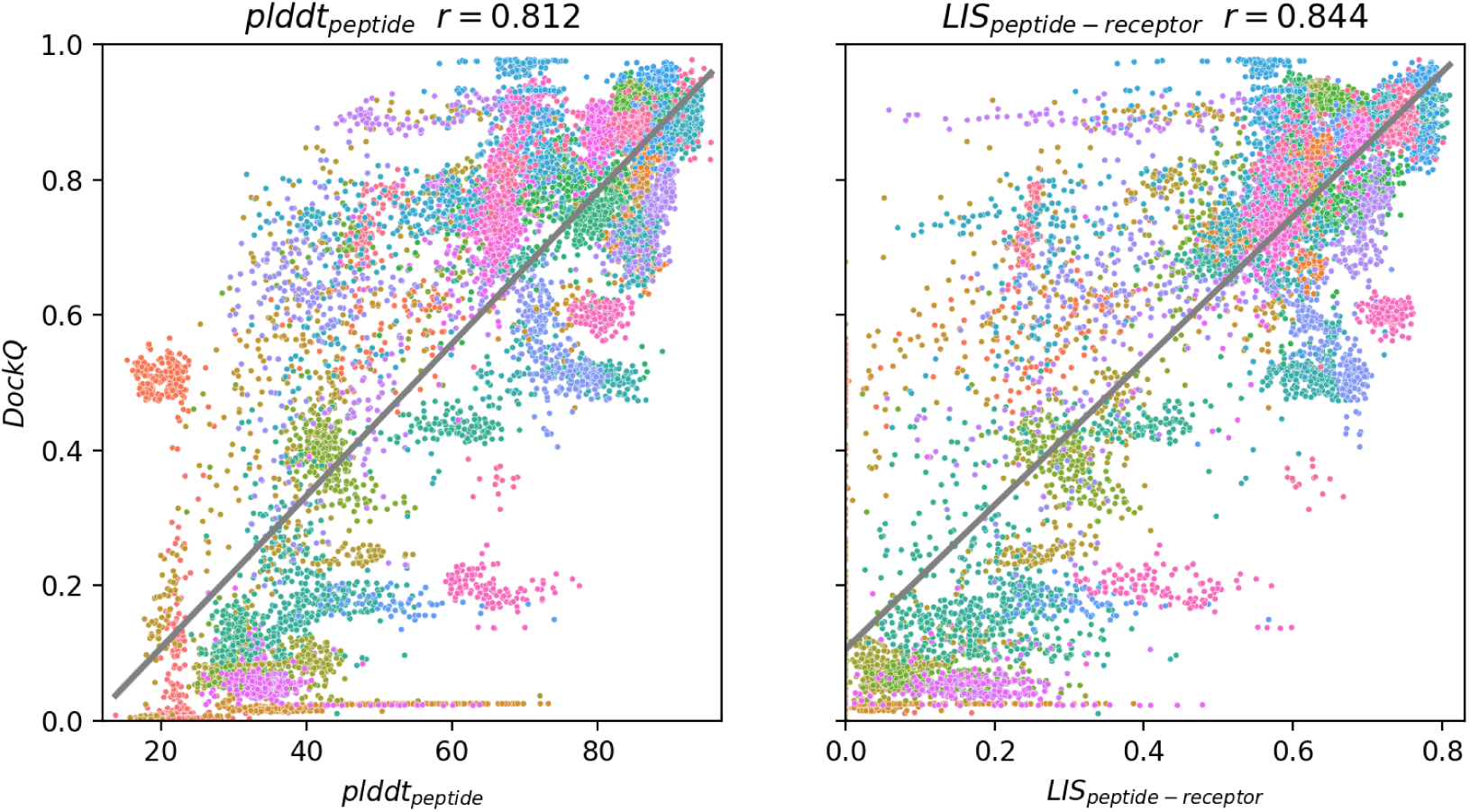
Correlation between DockQ and peptide-specific confidence scores. Scatter plots of DockQ against (left) peptide-restricted pLDDT (pLDDT_peptide_) and (right) the Local Interaction Score in the peptide-to-receptor direction (LIS_peptide_*_−_*_receptor_), computed across the pooled set of models generated for all 42 complexes with AlphaFold Multimer v2 and v3. Each point represents a single predicted model, colored by query (individual complex). The gray line shows the linear regression fit, with the corresponding Pearson correlation coefficient (r) reported in each panel title. Both scores show strong pooled correlation with DockQ, supporting their use as peptide-specific alternatives to global confidence metrics.

This discrepancy illustrates a critical caveat for the field: **correlations computed across heterogeneous complexes largely reflect the ability of a score to separate “easy” from “hard” targets, rather than its ability to rank models correctly within a given prediction ensemble, which is precisely the task required in practice, since the native structure is generally unavailable.** This limitation reinforces the need for scores (or combinations thereof) specifically calibrated for intra-target ranking of protein-peptide models rather than for cross-target quality estimation.

### IV. Practical Guidelines for Model Validation

#### IV.A. Combine Metrics

As shown above and in the literature, no single metric is sufficient for the comprehensive validation of a model.^75^ For multimers, examine pLDDT, PAE, pTM, and ipTM together. For interfaces, prioritize ipTM, and use pDockQ2 for ranking models. When disordered domains are included in the modeled protein, it is advisable to use scores that filter out the disordered regions or focus specifically on interface residues, such as actifpTM, ipSAE, or LIS-derived scores.

#### IV.B. Divide, conquer and confuse using allies !

- When modelling a complex, try modelling each chain individually. The presence or absence of a binding partner can induce conformational changes that may be revealed that way.
- When modelling a single chain, try modelling the probable IDRs and explicitly adding potassium cations as suggested above. This technique can allow for a fast assessment of whether the low pLDDT regions of a protein are actually disordered.
- Add ligands such as ions, small molecules or partner proteins to try to trigger conformational switches. This seems especially successful for nucleic acids, since our preliminary results suggest a sensitivity of the predicted conformation to ion type and number.

Overall, we recommend probing the modelled systems by exploiting the sensitivity of deep-learning schemes to noise in their input, a process which can be compared to some extent to scaling the temperature in a LLM.

#### IV.C. Visual Inspection

Sometimes, truth indeed is in the eye of the beholder. Beyond metrics and scores, visual inspection and human biochemical intuition paradoxically emerge as one of the most powerful approaches to critically assess the quality of a predicted structure. Inspecting the models “by eye” allows for the identification of problematic folds and steric clashes, as was evidenced by the prediction of one of the G-quadruplexes in this perspective (Figure 5, the severe superposition of potassium cations was identified by visual inspection). We recommend paying very close attention to regions with low pLDDT or high PAE. Tools such as ChimeraX,^88^ Pymol^89^ or PySSA^90^ can assist not only for the visualization of the predictions, but also provide automatic tools to identify clashes, unphysical geometries or inconsistent interfaces.

## Conclusion

In this perspective, we explored the new horizons for biomolecular structural predictions opened by the rise of deep-learning programs. Our assessment mainly focused on targets that remain difficult, such as transmembrane domains, nucleic acids and protein disorder. We focused on the possibilities and consequences of explicit ion modelling, showing that although AF3 predictions still lack reliability in their placement of ions, this explicit modelling can be hijacked in order to trigger new conformational predictions. We reviewed the available metrics to assess reliability in the context of AlphaFold predictions for both monomers and multimers, and provided general advice on how to critically determine whether a model can be trusted. We conclude given all the shortcomings listed in this perspective that while deeplearning methods constitute a progress in the field of structural biology, many limitations remain to be lifted.

## Material and methods

### MCTP/calcium predictions

For the MCTP4 protein, models were generated with AF3 both with and without calcium, ranging from 1 to 10 Ca^2+^ cations. APBS calculations were performed on the Poisson–Boltzmann web server (https://server.poissonboltzmann.org/). AlphaFill was used to identify calcium-binding sites according to experimental data available in the database.^35^ To identify experimentally determined structures containing C2 domains, we queried the RCSB Protein Data Bank using the Pfam annotation PF00168, corresponding to the canonical C2 domain family. All available structures associated with this Pfam entry were downloaded.

Calcium ions were then counted for each structure using a Python script, and the resulting distribution was plotted (see data in the Zenodo repository). For entries containing multiple structural models, only the first model was analyzed to avoid counting the same calcium ions multiple times.

### AlphaFold3 predictions, analysis and visualization

All AlphaFold3 predictions described in this paper were produced with the AlphaFold webserver with default parameters (https://alphafoldserver.com/).^4^ A single run of predictions was performed for each condition. The model ranked 0 by the server was deemed the most reliable and unless stated otherwise, single-structure predictions mentioned in the text refer to this model. RMSD alignments and analyses such as radius-of-gyration measurements were performed with the MDAnalysis software.^91^ Molecular visualization was performed with the VMD software.^92^

## Data availability

All AlphaFold predictions described in this paper are available as a Zenodo repository at the following link: https://zenodo.org/records/21534689

## Acknowledgement

The authors thank all the participants to the BeyondNobel2024 conference for their contribution to discussions. This work was supported by the ANR (MAGNETAU-ANR-21-CE29-0024, MITOFUSION-ANR-19-CE11-0018, DIVCON-ANR-21-CE13-0016, SUPERET-ANR-21-CE29-0013, PIRATE-ANR-21-CE45-0014) and by the “Initiative d’Excellence” program from the French State (Grant “DYNAMO”, ANR-11-LABX-0011-01 and grant “CACSICE”, ANR-11-EQPX-0008).

## Notes

https://zenodo.org/records/21534689

### Competing Interest Statement

The authors have declared no competing interest.

## References

(1) Rennie, M. L.; Oliver, M. R. Emerging frontiers in protein structure prediction following the AlphaFold revolution. Journal of the Royal Society Interface 2025, 22, 20240886.

(2) Graille, M.; Sacquin-Mora, S.; Taly, A. Best Practices of Using AI-Based Models in Crystallography and Their Impact in Structural Biology. Journal of Chemical Information and Modeling 2023, 63, 3637–3646.

(3) Versini, R.; Sritharan, S.; Aykac Fas, B.; Tubiana, T.; Aimeur, S. Z.; Henri, J.; Erard, M.; Nusse, O.; Andreani, J.; Baaden, M.; others A perspective on the prospective use of AI in protein structure prediction. Journal of chemical information and modeling 2024, 64, 26–41.

(4) Abramson, J. et al. Accurate structure prediction of biomolecular interactions with AlphaFold 3. Nature 2024, 630, 493–500.

(5) Peng, C.; Ni, W.; Liu, Q.; Hu, G.; Zheng, W. A comprehensive benchmarking of the AlphaFold3 for predicting biomacromolecules and their interactions. Briefings in Bioinformatics 2025, 26, bbaf616.

(6) Aimeur, S.; Fas, B. A.; Serfaty, X.; Santuz, H.; Sacquin-Mora, S.; Bizouarn, T.; Taly, A.; Baciou, L. Structural profiles of the full phagocyte NADPH oxidase unveiled by combining computational biology and experimental knowledge. J. Biol. Chem. 2024, 300, 107943.

(7) Brown, C. M.; Westendorp, M. S.; Zarmiento-Garcia, R.; Stevens, J. A.; Bruininks, B. M.; Rouse, S. L.; Marrink, S. J.; Wassenaar, T. A. An integrative modelling approach to the mitochondrial cristae. Communications Biology 2025, 8, 972.

(8) Nogales, E.; Mahamid, J. Bridging structural and cell biology with cryo-electron microscopy. Nature 2024, 628, 47–56.

(9) López-Sagaseta, J.; Urdiciain, A. Severe deviation in protein fold prediction by advanced AI: a case study. Scientific Reports 2025, 15, 4778.

(10) Škrinjar, P.; Eberhardt, J.; Studer, G.; Tauriello, G.; Schwede, T.; Durairaj, J. Evaluating generalization in protein–ligand cofolding methods. Nature Structural & Molecular Biology 2026, 33, 782–794.

(11) Chakravarty, D.; Lee, M.; Porter, L. L. Proteins with alternative folds reveal blind spots in AlphaFold-based protein structure prediction. Current Opinion in Structural Biology 2025, 90, 102973.

(12) Shin, O.-H.; Han, W.; Wang, Y.; Südhof, T. C. Evolutionarily Conserved Multiple C2 Domain Proteins with Two Transmembrane Regions (MCTPs) and Unusual Ca2+ Binding Properties*. 280, 1641–1651.

(13) Pucci, F.; Schug, A. Shedding light on the dark matter of the biomolecular structural universe: Progress in RNA 3D structure prediction. Methods 2019, 162, 68–73.

(14) Kagaya, Y.; Liu, B.; Kihara, D. Computational approaches for RNA structure prediction and design. Cell Reports Physical Science 2026, 7.

(15) Ochoa, S.; Milam, V. T. Direct Modeling of DNA and RNA Aptamers with AlphaFold 3: A Promising Tool for Predicting Aptamer Structures and Aptamer–Target Interactions. ACS Synthetic Biology 2025,

(16) Yu, H.; Qi, Y.; Ding, Y. Deep Learning in RNA Structure Studies. Frontiers in Molecular Biosciences 2022, 9, 869601.

(17) Bahai, A.; Kwoh, C. K.; Mu, Y.; Li, Y. Systematic benchmarking of deep-learning methods for tertiary RNA structure prediction. PLOS Computational Biology 2024, 20, e1012715.

(18) Justyna, M.; Antczak, M.; Szachniuk, M. Machine learning for RNA 2D structure prediction benchmarked on experimental data. Briefings in Bioinformatics 2023, 24, bbad153.

(19) Ludaic, M.; Elofsson, A. Limits of deep-learning-based RNA prediction methods. bioRxiv 2025, 2025.04.30.651414.

(20) Bernard, C.; Postic, G.; Ghannay, S.; Tahi, F. Has AlphaFold3 achieved success for RNA? Acta Crystallographica Section D 2025, 81, 49–62.

(21) Martinović, I.; Vlašić, T.; Li, Y.; Hooi, B.; Zhang, Y.; Šikić, M. A Comparative Review of Deep Learning Methods for RNA Tertiary Structure Prediction. bioRxiv 2024, 2024.11.27.625779.

(22) Nithin, C.; Kmiecik, S.; Błaszczyk, R.; Nowicka, J.; Tuszyńska, I. Comparative analysis of RNA 3D structure prediction methods: towards enhanced modeling of RNA–ligand interactions. Nucleic Acids Research 2024, 52, 7465–7486.

(23) Shen, T. et al. Accurate RNA 3D structure prediction using a language model-based deep learning approach. Nature Methods 2024, 1–12.

(24) Li, Y.; Feng, C.; Zhang, X.; Tsukiyama, S.; Feng, D.; Zhang, Y. DRfold2 Is a Deep Learning-Based Tool That Enables Efficient and Accurate RNA Structure Prediction. PLOS Biology 2026, 24, e3003659.

(25) Dégut, C.; Roovers, M.; Barraud, P.; Brachet, F.; Feller, A.; Larue, V.; Refaii, A. A.; Caillet, J.; Droogmans, L.; Tisné, C. Structural characterization of B. subtilis m1A22 tRNA methyltransferase TrmK: insights into tRNA recognition. Nucleic Acids Research 2019, 47, 4736–4750.

(26) Versini, R.; Baaden, M.; Bonvin, A. M.; Fuchs, P.; Taly, A. Full-Length Structural Modeling of Mitofusins with AlphaFold Reveals a Novel Cross-Type Dimerization and Insights into Oligomerization. bioRxiv 2026,

(27) Neupane, P.; Liu, J.; Cheng, J. Improving AlphaFold3 by Engineering MSA and Template Inputs. bioRxiv 2026,

(28) Cui, X.; Ge, L.; Chen, X.; Lv, Z.; Wang, S.; Zhou, X.; Zhang, G. Beyond static structures: protein dynamic conformations modeling in the post-AlphaFold era. Briefings in Bioinformatics 2025, 26, bbaf340.

(29) Seki, T.; Ohnuki, J.; Okazaki, K.-i. Conformational Dynamics of Na+-Pumping NADHQuinone Oxidoreductase during Na+ Translocation from AlphaFold-Facilitated Markov State Modeling. Journal of Chemical Information and Modeling 2026, 66, 4878–4887, PMID: 41930439.

(30) Sritharan, S.; Versini, R.; Petit, J. D.; Bayer, E. E.; Taly, A. Consensus structure prediction of A. thaliana’s MCTP4 structure using prediction tools and coarse grained simulations of transmembrane domain dynamics. PLOS ONE 2025, 20, 1–19.

(31) Genç, O.; Dickman, D. K.; Ma, W.; Tong, A.; Fetter, R. D.; Davis, G. W. MCTP is an ER-resident calcium sensor that stabilizes synaptic transmission and homeostatic plasticity. eLife 2017, 6, e22904.

(32) Téllez-Arreola, J. L.; Silva, M.; Martínez-Torres, A. MCTP-1 modulates neurotransmitter release in C. elegans. Molecular and Cellular Neuroscience 2020, 107, 103528.

(33) Liu, L.; Li, C.; Teo, Z. W. N.; Zhang, B.; Yu, H. The MCTP-SNARE Complex Regulates Florigen Transport in Arabidopsis. The Plant Cell 2019, 31, 2475–2490.

(34) Tuteja, N.; Mahajan, S. Calcium Signaling Network in Plants. Plant Signaling & Behavior 2007, 2, 79–85, PMID: 19516972.

(35) Hekkelman, M. L.; de Vries, I.; Joosten, R. P.; Perrakis, A. AlphaFill: enriching AlphaFold models with ligands and cofactors. Nature Methods 2023, 20, 205–213.

(36) Chakravarty, D.; Schafer, J. W.; Chen, E. A.; others AlphaFold predictions of fold-switched conformations are driven by structure memorization. Nature Communications 2024, 15, 7296.

(37) Chakravarty, D.; Porter, L. L. Fold-Switching Proteins. Annual Review of Biophysics 2026, 55, 17–37.

(38) del Alamo, D.; Sala, D.; Mchaourab, H. S.; Meiler, J. Sampling alternative conformational states of transporters and receptors with AlphaFold2. eLife 2022, 11, e75751.

(39) Wayment-Steele, H. K.; Ojoawo, A.; Otten, R.; Apitz, J. M.; Pitsawong, W.; Hömberger, M.; Ovchinnikov, S.; Colwell, L.; Kern, D. Predicting multiple conformations via sequence clustering and AlphaFold2. Nature 2024, 625, 832–839.

(40) Wolf-Watz, M.; Thai, V.; Henzler-Wildman, K.; Hadjipavlou, G.; Eisenmesser, E. Z.; Kern, D. Linkage between dynamics and catalysis in a thermophilic-mesophilic enzyme pair. Nature Structural & Molecular Biology 2004, 11, 945–949.

(41) Beckstein, O.; Denning, T. B., Elizabeth J. anlf Zipping and Unzipping of Adenylate Kinase: Atonsemble of Open^⇀^↽Closed Transitions. Journal of Molecular Biology 2009, 394, 160–176.

(42) Matsunaga, Y.; Fujisaki, H.; Terada, T.; Furuta, T.; Moritsugu, K.; Kidera, A. Minimum Free Energy Path of Ligand-Induced Trae. PLoS Computational Biology 2012, 8, e1002555.

(43) Westhof, E.; Sun, H.; Bu, F.; Miao, Z. The RNA-Puzzles Assessments of RNA-Only Targets in CASP16. *Proteins: Structure*, Function, and Bioinformatics 2025,

(44) Watkins, A. M.; Rangan, R.; Das, R. FARFAR2: Improved De Novo Rosetta Prediction of Complex Global RNA Folds. Structure 2020, 28, 963–976.e6.

(45) Townshend, R. J. L.; Eismann, S.; Watkins, A. M.; Rangan, R.; Karelina, M.; Das, R.; Dror, R. O. Geometric deep learning of RNA structure. Science 2021, 373, 1047–1051.

(46) Schneider, B.; Sweeney, B. A.; Bateman, A.; Cerny, J.; Zok, T.; Szachniuk, M. When will RNA get its AlphaFold moment? Nucleic Acids Research 2023, 51, 9522–9532.

(47) Xiao, B.; Shi, Y.; Huang, L. Enhancing RNA 3D structure prediction: A hybrid approach combining expert knowledge and computational tools in CASP16. Proteins 2026, 94, 230–238.

(48) Laine, E.; Grudinin, S.; Klypa, R.; Beauchêne, I. C. d. Navigating protein–nucleic acid sequence-structure landscapes with deep learning. Current Opinion in Structural Biology 2025, 95, 103162.

(49) Geng, Y.; Liu, C.; Miao, H.; Suen, M. C.; Xie, Y.; Zhang, B.; Han, W.; Wu, C.; Ren, H.; Chen, X.; Tai, H.-C.; Wang, Z.; Zhu, G.; Cai, Q. Crystal structures of distinct parallel and antiparallel DNA G-quadruplexes reveal structural polymorphism in C9orf72 G4C2 repeats. Nucleic Acids Research 2025, 53, gkaf879.

(50) Laneve, P.; Gioia, U.; Ragno, R.; Altieri, F.; Di Franco, C.; Santini, T.; Arceci, M.; Bozzoni, I.; Caffarelli, E. The Tumor Marker Human Placental Protein 11 Is an Endoribonuclease*. Journal of Biological Chemistry 2008, 283, 34712–34719.

(51) Malard, F.; Dias, K.; Baudy, M.; Thore, S.; Vialet, B.; Barthélémy, P.; Fribourg, S.; Karginov, F. V.; Campagne, S. Molecular basis for the calcium-dependent activation of the ribonuclease EndoU. Nature Communications 2025, 16, 3110.

(52) Renzi, F.; Caffarelli, E.; Laneve, P.; Bozzoni, I.; Brunori, M.; Vallone, B. The structure of the endoribonuclease XendoU: From small nucleolar RNA processing to severe acute respiratory syndrome coronavirus replication. Proceedings of the National Academy of Sciences 2006, 103, 12365–12370.

(53) McDonald, E. F.; Jones, T.; Plate, L.; Meiler, J.; Gulsevin, A. Benchmarking AlphaFold2 on peptide structure prediction. Structure 2023, 31, 111–119.e2.

(54) Alderson, T. R.; Pritišanac, I.; Ðesika Kolarić; Moses, A. M.; Forman-Kay, J. D. Systematic identification of conditionally folded intrinsically disordered regions by AlphaFold2. Proceedings of the National Academy of Sciences 2023, 120, e2304302120.

(55) Bret, H.; Gao, J.; Zea, D. J.; Andreani, J.; Guerois, R. From interaction networks to interfaces, scanning intrinsically disordered regions using AlphaFold2. Nature Communications 2024, 15, 597.

(56) Fayetorbay, R.; Timucin, A. C.; Timucin, E. Systematic Evaluation of AlphaFold2 and OpenFold3 on Protein-Peptide Complexes. bioRxiv 2026,

(57) Wang, Y.; Mandelkow, E. Tau in physiology and pathology. Nature Reviews Neuroscience 2016, 17, 22–35.

(58) Kellogg, E. H.; Hejab, N. M. A.; Poepsel, S.; Downing, K. H.; DiMaio, F.; Nogales, E. Near-atomic model of microtubule-tau interactions. Science 2018, 360, 1242–1246.

(59) Janke, C.; Magiera, M. M. The tubulin code and its role in controlling microtubule properties and functions. Nature Reviews Molecular Cell Biology 2020, 21, 307–326.

(60) Luo, Y.; Xiang, S.; Hooikaas, P. J.; van Bezouwen, L.; Jijumon, A. S.; Janke, C.; Förster, F.; Akhmanova, A.; Baldus, M. Direct observation of dynamic protein interactions involving human microtubules using solid-state NMR spectroscopy. Nature Communications 2020, 11, 18.

(61) Marien, J.; Prévost, C.; Sacquin-Mora, S. It Takes Tau to Tango: Investigating the Fuzzy Interaction between the R2-Repeat Domain and Tubulin C-Terminal Tails. Biochemistry 2023, 62, 2492–2502, PMID: 37499261.

(62) Shred, M.; Vangos, N. E.; Bayne, A. N.; Tetlalmatzi, S. C.; Peng, W.; Trempe, J.-F.; Sept, D.; Cianfrocco, M. A.; Brouhard, G. J. Conformational flexibility of tubulin dimers regulates the transitions of microtubule dynamic instability. bioRxiv 2025,

(63) Chakravarty, D.; Schafer, J. W.; Chen, E. A.; Thole, J. F.; Ronish, L. A.; Lee, M.; Porter, L. L. AlphaFold predictions of fold-switched conformations are driven by structure memorization. Nature communications 2024, 15, 7296.

(64) Škrinjar, P.; Eberhardt, J.; Durairaj, J.; Schwede, T. Have protein-ligand co-folding methods moved beyond memorisation? BioRxiv 2025, 2025–02.

(65) Makhatadze, G. I. The accuracy of electrostatic interactions captured by AI protein structure prediction models. Proceedings of the National Academy of Sciences 2026, 123, e2609610123.

(66) Chen, S.; Huh, J.; Sklar, C.; Gray, J. J. Predicting Supramolecular Self-Assembly of Peptide Structures with AlphaFold3. bioRxiv 2026,

(67) Chothia, C.; Lesk, A. M. The relation between the divergence of sequence and structure in proteins. The EMBO journal 1986, 5, 823–826.

(68) Olivella, M.; Gonzalez, A.; Pardo, L.; Deupi, X. Relation between sequence and structure in membrane proteins. Bioinformatics 2013, 29, 1589–1592.

(69) Jumper, J.; Evans, R.; Pritzel, A.; Green, T.; Figurnov, M.; Ronneberger, O.; Tunyasuvunakool, K.; Bates, R.; Žídek, A.; Potapenko, A.; others Highly accurate protein structure prediction with AlphaFold. Nature 2021, 596, 583–589.

(70) Baek, M. et al. Accurate prediction of protein structures and interactions using a three-track neural network. Science 2021, 373, 871–876.

(71) Akdel, M.; Pires, D. E.; Pardo, E. P.; Jänes, J.; Zalevsky, A. O.; Mészáros, B.; Bryant, P.; Good, L. L.; Laskowski, R. A.; Pozzati, G.; others A structural biology community assessment of AlphaFold2 applications. Nature Structural & Molecular Biology 2022, 29, 1056–1067.

(72) Wodak, S. J.; Vajda, S.; Lensink, M. F.; Kozakov, D.; Bates, P. A. Critical assessment of methods for predicting the 3D structure of proteins and protein complexes. Annual review of biophysics 2023, 52, 183–206.

(73) Yin, R.; Feng, B. Y.; Varshney, A.; Pierce, B. G. Benchmarking AlphaFold for protein complex modeling reveals accuracy determinants. Protein Science 2022, 31, e4379.

(74) Chungyoun, M.; Au, G.; Carpentier, B.; Puvada, S.; Thomas, C.; Gray, J. J. Deep Learning for Proteins Notebook Series Teaches AI for Biomolecular Structure Prediction and Design. The Biophysicist 2026, 1.

(75) Genz, L. R.; Nair, S.; Nagar, N.; Topf, M. Assessing scoring metrics for AlphaFold2 and AlphaFold3 protein complex predictions. Protein Science 2025, 34, e70327.

(76) Zhu, W.; Shenoy, A.; Kundrotas, P.; Elofsson, A. Evaluation of AlphaFold-Multimer prediction on multi-chain protein complexes. Bioinformatics 2023, 39, btad424.

(77) Varga, J. K.; Ovchinnikov, S.; Schueler-Furman, O. actifpTM: a refined confidence metric of AlphaFold2 predictions involving flexible regions. Bioinformatics 2025, 41, btaf107.

(78) Kim, A.-R.; Hu, Y.; Comjean, A.; Rodiger, J.; Mohr, S. E.; Perrimon, N. Enhanced Protein-Protein Interaction Discovery via AlphaFold-Multimer. bioRxiv 2024,

(79) Kim, A.-R.; Comjean, A.; Veal, A.; Rodiger, J.; Han, M.; Hu, Y.; Perrimon, N. FlyPredictome: A structural atlas of predicted protein-protein interactions in Drosophila. bioRxiv 2026,

(80) Dunbrack, R. L. Res ipSAE loquuntur: What’s wrong with AlphaFold’s ipTM score and how to fix it. bioRxiv 2025,

(81) Mirdita, M.; Schütze, K.; Moriwaki, Y.; Heo, L.; Ovchinnikov, S.; Steinegger, M. ColabFold: making protein folding accessible to all. Nature Methods 2022, 19, 679–682.

(82) Genz, L. R.; Mulvaney, T.; Nair, S.; Topf, M. PICKLUSTER: a protein-interface clustering and analysis plug-in for UCSF ChimeraX. Bioinformatics 2023, 39, btad629.

(83) Reguei, A.; Murail, S. Af-analysis: a Python package for Alphafold analysis. Journal of Open Source Software 2025, 10, 7577.

(84) Álvarez-Salmoral, D.; Borza, R.; Maiella, C.; Kwee, B. P.; Xie, R.; Joosten, R. P.; Hekkelman, M. L.; Perrakis, A. AlphaBridge: tools for the analysis of predicted biomolecular complexes. bioRxiv 2025,

(85) Fadini, A. et al. Highlights of Model Quality Assessment in CASP16. *Proteins: Structure*, Function, and Bioinformatics 2026, 94, 314–329.

(86) Evans, R. et al. Protein complex prediction with AlphaFold-Multimer. bioRxiv 2022,

(87) Mirabello, C.; Wallner, B. DockQ v2: improved automatic quality measure for protein multimers, nucleic acids, and small molecules. Bioinformatics 2024, 40, btae586.

(88) Meng, E. C.; Goddard, T. D.; Pettersen, E. F.; Couch, G. S.; Pearson, Z. J.; Morris, J. H.; Ferrin, T. E. UCSF ChimeraX: Tools for structure building and analysis. Protein Science 2023, 32, e4792.

(89) Schrödinger, LLC

(90) Kullik, H.; Urban, M.; Schaub, J.; Loidl-Stahlhofen, A.; Zielesny, A. PySSA for Windows: End-User Protein Structure Prediction and Visual Analysis with ColabFold and PyMOL. Journal of Chemical Information and Modeling 2025, 65, 5839–5846, PMID: 40458895.

(91) Michaud-Agrawal, N.; Denning, E. J.; Woolf, T. B.; Beckstein, O. MDAnalysis: A toolkit for the analysis of molecular dynamics simulations. Journal of Computational Chemistry 2011, 32, 2319–2327.

(92) Humphrey, W.; Dalke, A.; Schulten, K. VMD – Visual Molecular Dynamics. Journal of Molecular Graphics 1996, 14, 33–38.

